# Robust Acquisition of High Resolution Spatial Transcriptomes from Preserved Tissues with Immunofluorescence Based Laser Capture Microdissection

**DOI:** 10.1101/2021.07.13.452123

**Authors:** Xiaodan Zhang, Chuansheng Hu, Chen Huang, Ying Wei, Xiaowei Li, Miaomiao Hu, Hua Li, Ji Wu, Daniel M. Czajkowsky, Yan Guo, Zhifeng Shao

## Abstract

The functioning of tissues is fundamentally dependent upon not only the phenotypes of the constituent cells but also their spatial organization in the tissue. However, acquisition of comprehensive transcriptomes of spatially- and phenotypically-defined cells *in situ* remains challenging. Here we present a general and robust method based on immunofluorescence-guided laser capture microdissection (immuno-LCM-RNAseq) to acquire finely resolved spatial transcriptomes including isoforms with as few as tens of cells from snap-frozen or RNAlater-treated clinical tissues, circumventing the problem of significant RNA degradation during this time-consuming process. The efficacy of this approach is exemplified by the characterization of differences at the transcript isoform level between the mouse small intestine lacteal cells at the tip versus the main capillary body. With the extensive repertoire of specific antibodies that are presently available, our method provides a powerful means by which spatially resolved cellular states can now be delineated *in situ* with preserved tissues. Moreover, such high quality spatial transcriptomes defined by immunomarkers can be used to compare with the phenotypes derived from single-cell RNAseq of dissociated cells as well as applied to bead-based spatial transcriptomic approaches that require such information *a priori* for cell identification.

## Introduction

It is now well-recognized that the functioning of any tissue, whether healthy or diseased, is uniquely determined by the diverse constituent cells interacting within a highly structured three-dimensional architecture. Thus, techniques that can characterize the transcriptomes of defined cells within a tissue whilst retaining their spatial information are of immense interest (Crosetto et al. 2005; Moor and Itzkovitz 2017). Recent developments to this end can be generally divided into two broad categories: methods based on spatially resolved imaging of fluorescence *in situ* hybridization (FISH) and those based on spatial profiling of transcripts using next generation sequencing. For the former, while directly imaging the FISH signals provides a straightforward means of detection and quantification, the transcriptomes determined by these methods only include those transcripts that are fully annotated and for which suitable probes can be designed, which limits the detectable transcripts, particularly those from novel genes (Asp et al. 2020; Liao at al. 2021). This issue can be avoided with sequencing-based approaches, such as Slide-seq (Rodriques et al. 2019) and HDST (Vickovic et al. 2019) that are both based on the sequencing of transcripts captured on tagged beads. Yet, identifying individual cells in the tissue that are associated with the bead-bound transcripts is less straightforward and often requires knowledge of the transcriptomes of the constituent cells in the tissue *a priori*, ideally acquired under the same (physiological) conditions. Moreover, owing to the enrichment at the 3’ end of the transcripts, these sequencing-based methods do not resolve transcript isoforms, which, it should be noted, is also a problem with the imaging-based approaches as well.

In this regard, the more traditional laser capture microdissection (LCM) (Emmert-Buck et al. 1996) combined with RNAseq is a powerful approach with its own unique strengths for the acquisition of spatial transcriptomes, since cells are directly selected from the tissue with complete knowledge of their spatial location for an unbiased delineation of their transcriptome. Although identifying cells for LCM based on cell morphology alone is possible in some cases (Hawrylycz et al. 2012; Peng et al. 2016; Murray 2018), the vast majority of cell types in a tissue cannot be distinguished by morphology alone (Hanahan and Weinberg 2011). Therefore, use of specific phenotype markers, especially immunofluorescence (IF)-based cell type identification, remains the most attractive strategy to acquire comprehensive transcriptomes *in situ* with LCM (Fend et al. 1999; Murakami et al. 2000). To date, however, despite some commonly-held perceptions, there are, in fact, only limited successes of highly effective integration of IF with LCM for transcriptome acquisition (Liao et al. 2021; Murray 2018; Rao et al. 2021; Tangrea et al. 2011). This difficulty is primarily owing to the fact that RNA is often significantly degraded during IF labelling as a result of the activities of the intrinsic and ubiquitous exogenous RNases, even with the use of potent recombinant RNase inhibitors (Feng et al. 1999; Murakami et al. 2000; Tangrea et al. 2011). Several strategies to overcome this problem have been proposed, including drastically shortening the incubation time during labelling while using much higher concentrations of antibodies, or using high salt solutions to reduce the RNases activities (Brown and Smith 2009; Grimm et al. 2004; Tangrea et al. 2011;Nichterwitz et al. 2016; Baccin et al. 2020). However, neither method has proven sufficiently robust with moderate-to-high RNase-content tissues in terms of both high quality imaging and high RNA quality to enable comprehensive cell-type specific transcriptomic analyses.

In this paper, we present a broadly applicable immuno-LCM-RNAseq method that enables high quality RNAseq with a wide range of tissues for spatial transcriptome acquisition from sections of either snap-frozen or RNAlater preserved clinical tissues. The power of this approach is demonstrated here with an initial characterization of the transcriptomic differences between the lacteal tip and main body cells in the mouse small intestine, which implicate several genes that may play critical roles in the maintenance and morphology of this fine structure. We anticipate that this method will become indispensable in the acquisition of spatial transcriptomes of phenotype-defined cells in their native environment, especially with preserved tissues of clinical significance.

## Results

### Evaluation of the effectiveness of the present immunolabelling methods for LCM-seq

We first re-evaluated the image quality and extent of RNA protection of the existing approaches used for immuno-LCM, namely “Rapid Immunostaining” (hereafter called the “Rapid protocol”) (Nichterwitz et al. 2016) and using high concentrations of salt (the “high-salt protocol”) (Brown and Smith 2009), during immunostaining with snap-frozen tissues. The Rapid protocol entails the incubation of the sections with high concentrations of antibodies (1:10 to 1:25 dilutions, about ten-fold greater than that recommended for regular IF staining) in the presence of high concentrations of a potent RNase inhibitor (1 – 2 U/μl) (Grimm et al. 2004). The complete procedure, which typically requires a minimum of several hours with conventional IF staining (Im et al. 2004), is drastically shortened to only 3 to 15 minutes. The high-salt protocol involves the addition of 2 M NaCl to the antibody incubation solution but otherwise follows the conventional several-hour procedure to complete. Neither of these protocols recommended a pre-block step although such pre-blocking is known to reduce background in most immunolabelling procedures to improve image quality. Closely following these protocols, we examined their effectiveness to produce high quality fluorescence images and to preserve RNA quality using an anti-Lyve1 antibody (1:25 dilution, 40 μg/ml) to label lymphatic vessels in frozen sections of the mouse brain and small intestine. RNA quality was assessed with entire sections after the immunostaining step but without laser dissection. For the brain sections, which are largely devoid of endogenous RNases, both methods yielded good quality RNA with RIN values of ~9.0 (Fig. 1A-F and Supplemental Fig. S1). However, the contrast (S/N) in the fluorescent images obtained with either method (using 15 min incubation for the Rapid protocol) was poor when compared to that obtained with the conventional protocol (Fig. 1G-L and Supplemental Fig. S1). In fact, such low quality images make unambiguous identification of targeted cells, a prerequisite for LCM, nearly impossible.

**Figure 1.**
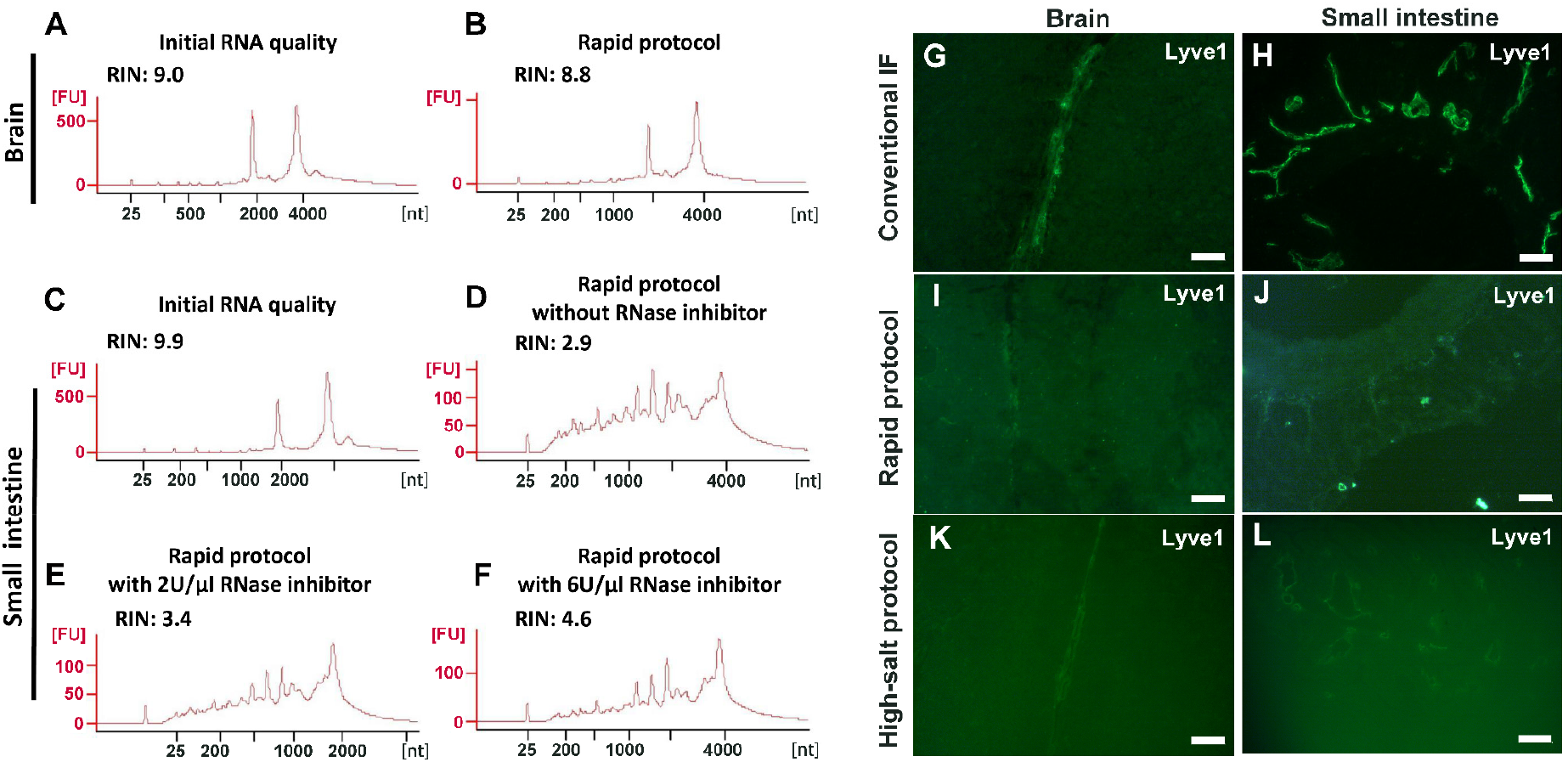
Evaluation of the Rapid and high-salt immunofluorescence staining protocols with frozen sections of the mouse brain and small intestine. (**A-F**) RNA quality assessment for frozen sections after immunofluorescence labelling using the Rapid protocol. For the brain sections, the RNA quality after rapid immunolabelling (B) is comparable to the initial quality (A). For the small intestine with a high endogenous RNase content, the RNA were severely degraded (D) even when high concentrations of RNase inhibitor were present (E-F). (**G-L**) Quality of IF images with the mouse brain (G, I, K) or the small intestine (H, J, L). Here, anti-Lyve1 was used to identify lymphatic vessels. With both protocols (I-L), the image quality in both tissues is poor when compared with the control (G, H). Scale bar: 50 μm (**G, I, K, L**) and 100 μm (**H, J**).

For the mouse small intestine, which has a much higher RNase content than the brain (Uhlén et al. 2015; Walker et al. 2016; https://www.proteinatlas.org/ENSG00000129538-RNASE1/tissue), only the high-salt protocol provided good RNA protection, albeit with a moderate degradation with increased incubation time (RIN: 8.6 at 5 hrs vs 7.0 overnight) (Supplemental Fig. S1). However, with this approach, not only was the IF image poor but the tissue structure was severely compromised (Fig. 1L, Supplemental Fig. S1). This structural degradation essentially prohibited any meaningful investigation of the spatial transcriptome as the section had lost many of its constituent cells. For the Rapid protocol, the procedure failed to provide effective RNA protection and the extracted RNA only had a RIN of 2.9. At the same time, the fluorescence image remained poor (Fig. 1J). We note that this degradation of RNA with the Rapid protocol occurred despite the presence of recombinant RNase inhibitors (2 U/μl). With a further increase of the RNase inhibitors up to 6 U/μl (3-fold higher than the recommended value), the observed improvements were disappointing if at all (Fig. 1E-F).

One possible reason for this failure of the RNA inhibitors to protect RNA in RNase-rich sections is that the diffusion of this inhibitor, a relatively large protein (~50 KD with an estimated dimension of 7 x 6 x 3 nm^3^ (ref. Hofsteenge 1997)) (Supplemental Fig. S2), might be too slow in tissue sections, as tissues are essentially dense, gel-like matrices (Davies et al. 2002). This might be a problem further exacerbated by the required dehydration-rehydration steps. Hence, it could be that it takes too long for the inhibitors to reach deep inside the section following rehydration, leading to rapid RNA degradation by the abundant endogenous RNases before the inhibitors arrive. As such, this type of RNase inhibitor is poorly suited for use with tissue sections.

### Development of a strategy using a small molecule RNase inhibitor for generally effective immuno-LCM-RNAseq

Given the ineffectiveness of the recombinant RNase inhibitors, we reasoned that small molecule RNase inhibitors might confer a much better protection owing to their faster diffusion within the rehydrated tissue section and thus quicker deactivation of the endogenous RNases. In this regard, it has been known that nucleoside analogues are potent inhibitors of many classes of nucleases (Russo et al. 2001). Among the many candidates, the ribonucleoside-vanadyl complex (RVC) (Berger and Birkenmeier 1979) is particularly attractive since it is a transition-state analogue (Lienhard et al. 1972; Leon-Lai et al. 1996): these complexes specifically bind to the catalytic site of ribonuclease (Supplemental Fig. S2) and should be broadly effective against many different RNases (Berger and Birkenmeier 1979). Although these complexes have been used to preserve RNA in tissues in a few studies previously (Shieh et al. 2018; Credle et al. 2017; Lyubimova et al. 2013), they have been largely superseded by the more potent recombinant RNA inhibitors in most other experiments (Shapiro 2001; Dickson et al. 2005; Mita et al. 2021; Popella et al. 2021; Qian et al. 2020; Arizti-Sanz et al. 2020). Whether RVC could provide robust RNA protection following immunolabelling, or immuno-LCM, which typically require many hours to complete, has not been previously examined. To this end, to compare its effectiveness with the aforementioned approaches, we first examined mouse brain frozen sections during the standard long time immunostaining with the anti-Lyve1 antibody that identifies lymph vessels with different concentrations of RVC in the incubating solutions. As with the standard immunofluorescence staining, cooled acetone fixed brain sections were first blocked (to reduce non-specific binding) for 15 minutes, followed by incubation with the primary antibody (1:100 dilution as recommended; ~10 μg/ml) for 3.5 hours, followed by secondary antibody (10 μg/ml) incubation for 1 hour at 4°C. We compared the results in 3 different RVC concentrations in all of the buffer solutions: 2.5 mM, 5 mM or 10 mM. We found that RVC had negligible effects on antigen-antibody interactions at these concentrations. As shown in Supplemental Fig. S3, the resultant IF images were of high contrast, essentially the same as those without the addition of any RVC. When the RNA quality of these treated sections was assessed, we found that samples with 5 mM or 10 mM RVC provided superb RNA protection with RIN > 9.5. But at 2.5 mM RVC, some RNA degradation was apparent (RIN 7.2) (Supplemental Fig. S4). These results demonstrate that, at least for brain sections, a minimum of 5 mM RVC is required for conventional (high quality) immunostaining to ensure fully protected RNA. We should also indicate that, in the process of performing these experiments, we found that the potency of the RVC solutions to protect RNA decreased over the course of days (similar to previous reports) (Berger and Birkenmeier 1979). However, use of freshly prepared RVC solutions proved to be a simple, effective remedy of this problem.

We next examined the effectiveness of RVC with other frozen tissue sections containing moderate-to-high levels of RNase (Uhlén et al. 2015; Walker et al. 2016; https://www.proteinatlas.org/ENSG00000129538-RNASE1/tissue). With the mouse small intestine, we found that 5 mM RVC was not sufficient to fully protect the RNA (RIN 6.3), but with 10 mM RVC, the quality of RNA was significantly improved (RIN 8.1) while the IF images remained excellent (Supplemental Fig. S3 – S4). Further increasing the RVC concentration (up to 20 mM) resulted in a moderate improvement in RNA quality (RIN 8.8) but with some adverse effects on the antibody-antigen interactions, leading to a deterioration of the IF images (Supplemental Fig. S3 - S4).

Therefore, we used 10 mM RVC as the optimal working concentration to examine its protective effect with various mouse frozen tissue sections: the stomach, liver, kidney, colon, spleen and testis (Fig. 2). For each of these tissues, and for all antibodies tested so far, 10 mM RVC invariably provided robust RNA protection with RIN values ranging from 7.3 (spleen) to 9.7 (stomach) (Fig. 2). When RVC was absent, the best RNA quality that could be obtained with these tissues was around RIN 3.2 (testis), far below that required for high quality RNA profiling. Similar to the mouse brain sections, high quality IF images were obtained with all of these tissues at 10 mM RVC (Fig. 3). Such a high quality is more than adequate for precise microdissection.

**Figure 2.**
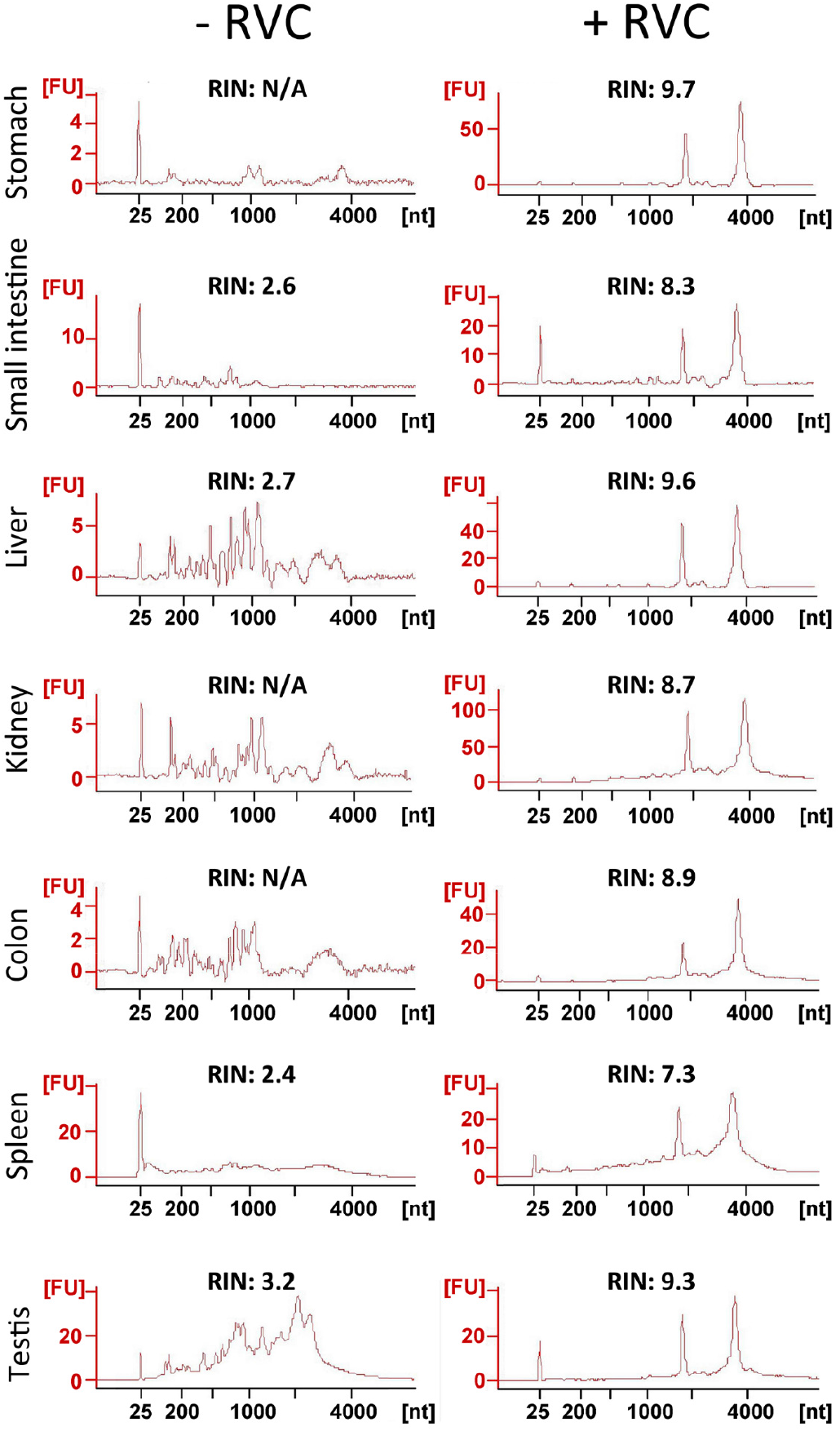
Assessment of RNA quality from sections of various snap-frozen mouse tissues after the standard immunolabelling procedure (details in the text) in the presence or absence of 10 mM RVC. As shown by the RIN values, RVC provided superb RNA protection in all of these RNase-rich tissues during the lengthy immunolabelling procedure.

**Figure 3.**
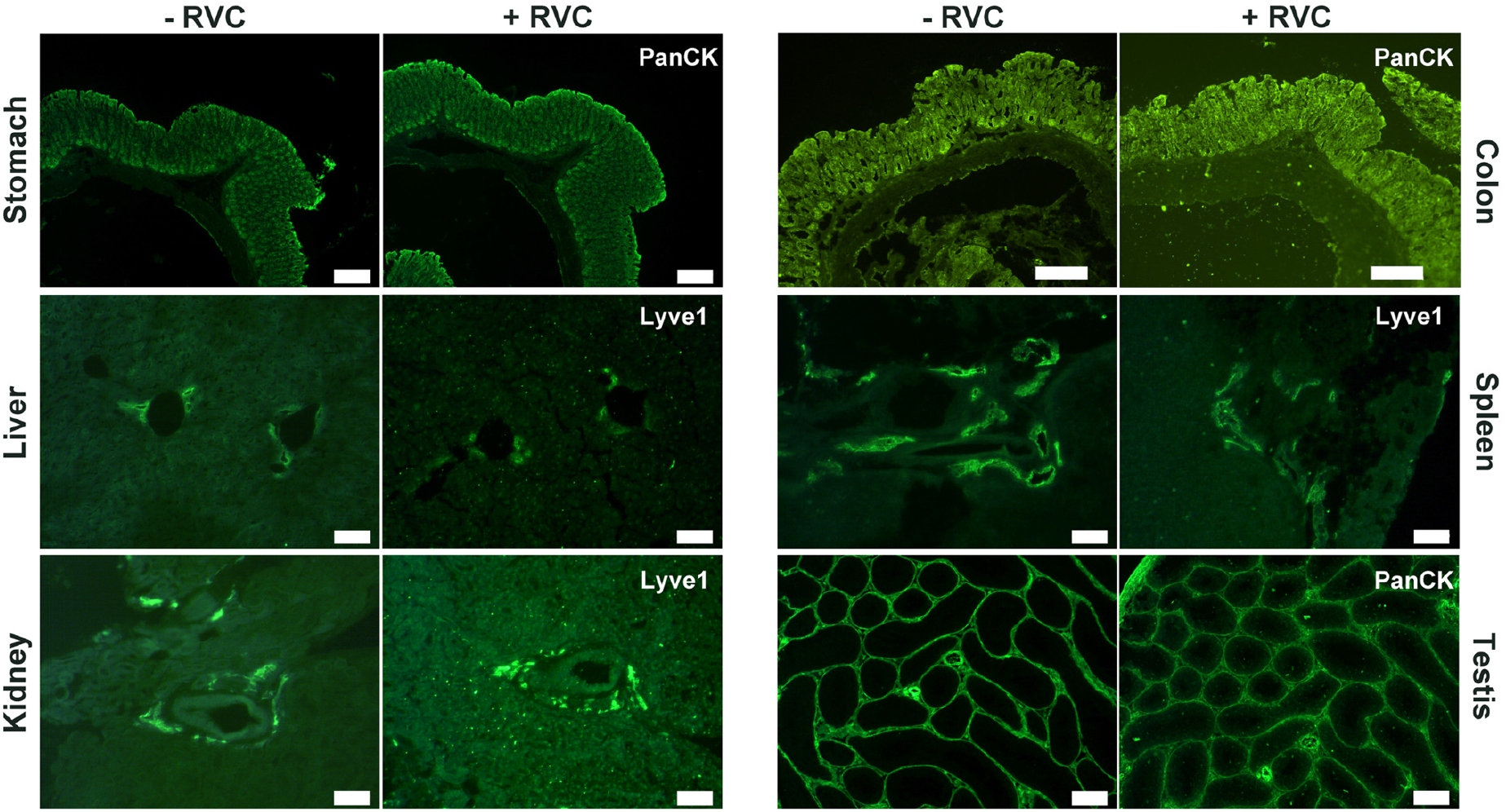
High quality IF images obtained from various snap-frozen tissue sections with standard immunolabelling procedures in the presence of 10 mM RVC. The presence of RVC in the incubation solution apparently has no adverse effect on antibody-antigen interactions. Here, anti-Lyve1 labels lymphatic vessels and anti-PanCK labels epithelial cells. Scale bar: 200 μm (stomach, colon, testis), 100 μm (liver, kidney, spleen)

Based on the above findings, we finally examined the efficacy of the complete LCM-RNAseq procedure with IF-based cell identification, incorporating 10 mM RVC in all incubation/wash steps (immuno-LCM-RNAseq; see Fig. 4 for the protocol flow chart). From sections of the mouse small intestine, the PanCK antibody was used to identify epithelial cells (Fig. 5A, upper panel). The identified target cells were then manually marked and automatically laser dissected and collected with the Zeiss PALM MicroBeam LCM system. The dissection and collection process required 45~60 minutes to complete and was performed at room temperature. As a preliminary test, we microdissected about 2,300 cytokeratin-positive epithelial cells (Ep2300) from the crypt region (within 100 μm from the submucosa) of 12 μm thick sections of the small intestine (Fig. 5A, Supplemental Fig. S5-S6, Supplemental Table S1). Examining material leftover after the laser microdissection, we found that the RNA integrity was sufficiently retained (RIN 8.3) (Fig. 5B). Since the dissection was performed under ambient conditions in open air where a small amount of water is condensed on hydrophilic surfaces (Clement-Ziza et al. 2008), such robust RNA protection suggests that the residual RVC after section dehydration (as necessary for laser microdissection) remained effective against airborne RNase contaminants (Clement-Ziza et al. 2008).

**Figure 4.**
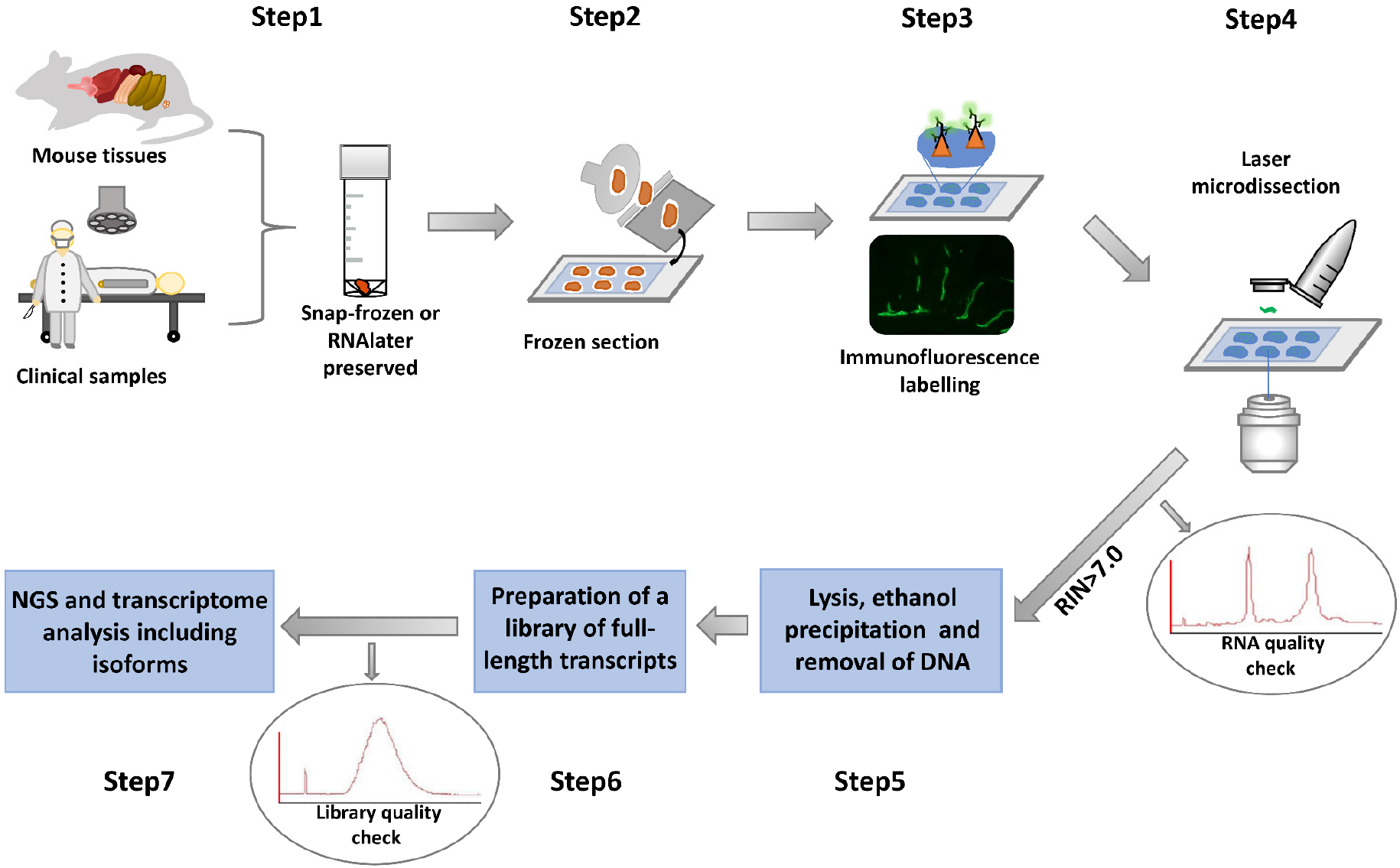
Overview of the immuno-LCM-RNAseq method. Either snap-frozen or RNAlater-preserved tissues is sectioned in a cryostat at appropriate temperatures. After the immunofluorescence guided laser microdissection, it is critical to use the leftover materials for an RNA quality check. Only with a high RIN value should the experiment continue. Although the cDNA library could be prepared with many methods, the Ovation SoLo system performed well and allowed for the analysis of full transcripts and isoforms with high fidelity. Before sequencing, the quality of the library should be checked with a 2100 Bioanalyzer to ensure the quality of the final result.

**Figure 5.**
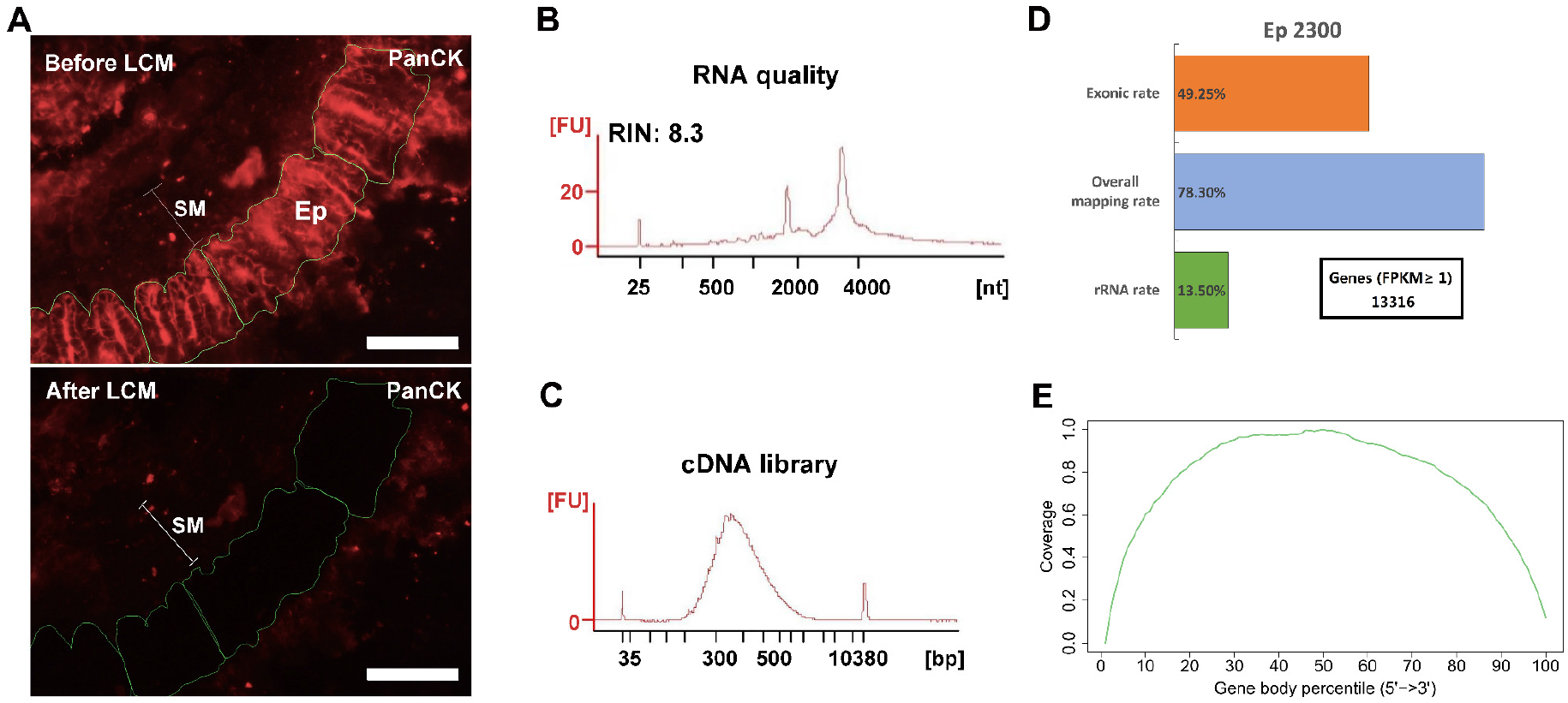
Immumo-LCM-RNAseq with ~2300 cytokeratin positive epithelial cells (Ep2300). (**A**) A typical anti-PanCK stained fluorescence section before (upper) and after (lower) laser microdissection. SM: submucosal layer; Ep: cytokeratin positive cells. Scale bar: 100 μm. An image showing the nuclei stained with Hoechst is shown in Supplemental Fig. S6. (**B**) High RNA quality of the leftover materials after microdissection. (**C**) cDNA library quality assessment. (B) and (C) were obtained with the 2100 Bioanalyzer. (**D**) Summary of the sequencing data quality with more than 13,000 expressed genes detected. (**E**) As expected from the Ovation SoLo system, normal reads coverage over the transcripts was obtained without 3’ enrichment. This coverage is required for full transcript and isoform analysis.

One problem with the use of RVC is its effect on downstream procedures, including reverse transcription and/or PCR that are required for the construction of cDNA sequencing libraries, owing to its adverse effect on polymerases (Lau et al. 1993) (Step 6, Fig. 4). Hence, RVC must be removed from the dissected materials before the downstream procedures can be performed properly. We used the ethanol precipitation/extraction method to purify RNA for the downstream construction of the cDNA library (removing residual DNA by DNase digestion) in this protocol. For the Ovation SoLo RNAseq system, which is optimized for low input RNA down to 10 pg and also adequate for mRNA transcript isoform analysis, the purified RNA was sufficient to construct the cDNA library (Nguyen et al. 2018). As shown in Fig. 5C, the cDNA library obtained is indeed of high quality with a proper fragment distribution. After sequencing, we obtained about 16 million clean reads for this sample (Supplemental Table S2). The overall mapping rate was 78% and the exonic rate was 49%, both consistent with the expected outcomes of the Ovation SoLo system (Fig. 5D). Using FPKM ≥ 1 as the threshold, more than 13,000 expressed genes were identified (Fig. 5D). The reads coverage across the gene body showed no bias towards the 5’ or 3’ end (Fig. 5E), an indication of high quality RNA (Wang et al. 2016). Such a quality should be appropriate to investigate full-length transcripts and isoforms. Thus, these results demonstrate the effectiveness of our immuno-LCM-RNAseq protocol to obtain high quality transcriptomes (Fig. 4).

### Determining the working limit of immuno-LCM-RNAseq

Since any RNA extraction procedure will lead to a certain amount of material loss, we sought to determine the minimal number of dissected cells required for successful immuno-LCM-RNAseq. Again, using the PanCK antibody to identify the epithelial cells in the snap-frozen mouse small intestine sections, we laser dissected 630, 230, or 63 cells, also in the crypt region. Together with the Ep2300, we evaluated the consistency between the obtained transcriptomes of these samples (Fig. 6) at a comparable level of sequencing depth (13 – 20 million clean reads) (Supplemental Table S2). Despite the significant difference in the amount of material collected (> 30 fold), a similar number of expressed genes were identified under the same threshold (FPKM ≥ 1) (Fig. 6C), demonstrating a high efficiency of this method. The reads coverage across the gene body also shows no bias towards the 5’ or 3’ end (Fig. 6B). However, we noted that at the low end of ~ 60 cells, the number of detected genes is slightly lower than the other samples, most likely owing to material loss in the RNA purification step. Nonetheless, the Pearson correlation (R) among these samples remained high (1 kb bin) (average R = 0.88) (Fig. 6D), demonstrating the reproducibility over a large range of collected input materials. Therefore, our current protocol with ethanol precipitation combined with the Ovation SoLo system remains robust down to a few tens of dissected cells.

**Figure 6.**
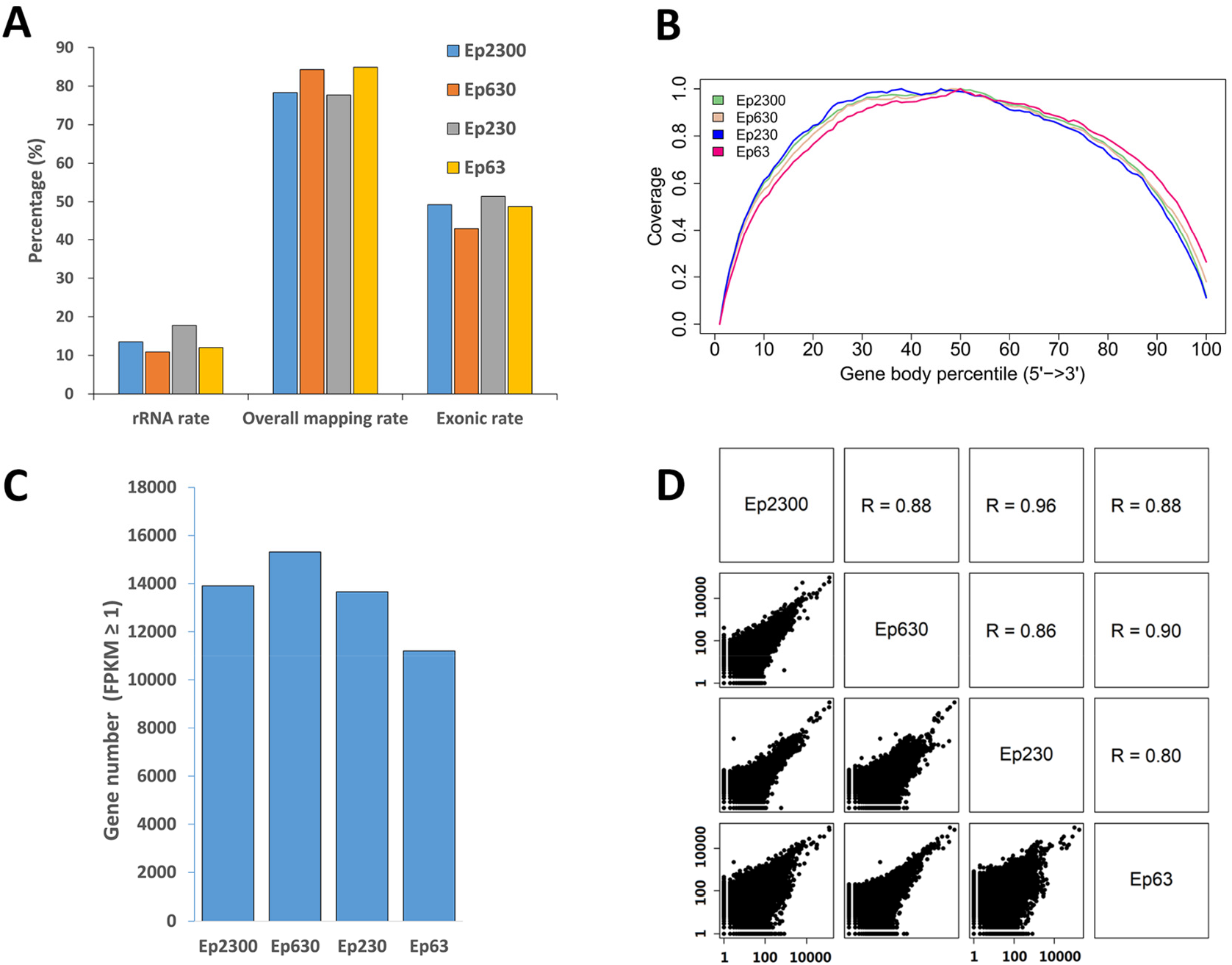
Assessment of the data quality and reproducibility with different amounts of input material dissected from the mouse small intestine using PanCK as the marker. The number of cells in each sample is indicated by the value following “Ep”, i.e., Ep63 = 63 cells. (**A**) Proportion of the overall mapping rate, the rRNA mapping rate and exonic rate of the samples. Despite a greater than 30-fold difference in the input material, the data quality remained consistent. (**B**) Reads distribution over the gene body is normal, without 5’ or 3’ end enrichment. (**C**) The number of expressed genes detected in all of the samples is similar, but is a little lower with the Ep63 sample, suggesting that the RNA extraction step could be further optimized. (**D**) Pairwise scatter plots between all samples. These Pearson correlation coefficients indicate that these results are consistent with each other.

This analysis also enables a comparison with results obtained in a recent LCM-RNAseq study of bone marrow tissue (that has a similar level of RNase content as the small intestine (Uhlén et al. 2015; https://www.proteinatlas.org/ENSG00000129538-RNASE1/tissue; https://www.proteinatlas.org/ENSG00000169385-RNASE2/tissue)) in which the Rapid protocol was used during immuno-labelling. Although the main focus of this work was the characterization of the transcriptomic details of mixtures of 200-300 cells in different niches, and not a comprehensive understanding of phenotype-specific cells, their results serve a useful indicator of what can be presently achieved with the Rapid protocol. From a comparable number of cells, and at a similar sequencing depth, we were able to identify more genes (13521 vs 3266), with a lower rRNA mapping rate (14% vs 65%) and a lower intergenic mapping rate (7% vs 15%), as well as superior overall and exonic mapping rates (Supplemental Table S2), with our approach. Thus, while this Rapid protocol method can provide useful information about the cellular organization of tissues, our method enables a more comprehensive understanding of the spatial transcriptome.

### Spatially defined expression: comparing the cells at the tip and in the main capillary body of the lacteal

We next applied immuno-LCM-RNAseq to examine an interesting fine structure, namely, the mouse small intestine lacteal, which cannot be readily resolved based on cell morphology alone (Supplemental Fig. S7). The lacteal is the lymphatic capillary in the small intestine villi with crucial roles in fat absorption and gut immune response (Bernier-Latmani and Petrova 2017). However, unlike other lymphatic vessels, the lacteal cells are found to be moderately proliferative and exhibit long filopodia-like protrusions at the lacteal end (the “tip” cells) (Bernier-Latmani and Petrova 2017). As shown in Fig. 7A, these fine capillaries can be clearly identified with the anti-Lyve1 antibody. We first dissected and collected ~150 Lyve1 positive cells from the main body of the lacteal (50 – 70 μm away from the lacteal tip, referred to as the “tube”) (Fig. 7A-B, Supplemental Table S2 - S3) with excellent RNA quality (RIN 8.7, Supplemental Fig. S8). With a total of 4 replicates, we obtained 49 – 63 million clean reads for each sample and the average overall mapping rate was 80%. The average Pearson correlation coefficient between all pairwise comparisons was 0.86. On average, more than 11,000 expressed genes were detected in each sample with FPKM ≥ 1 (Supplemental Table S2).

**Figure 7.**
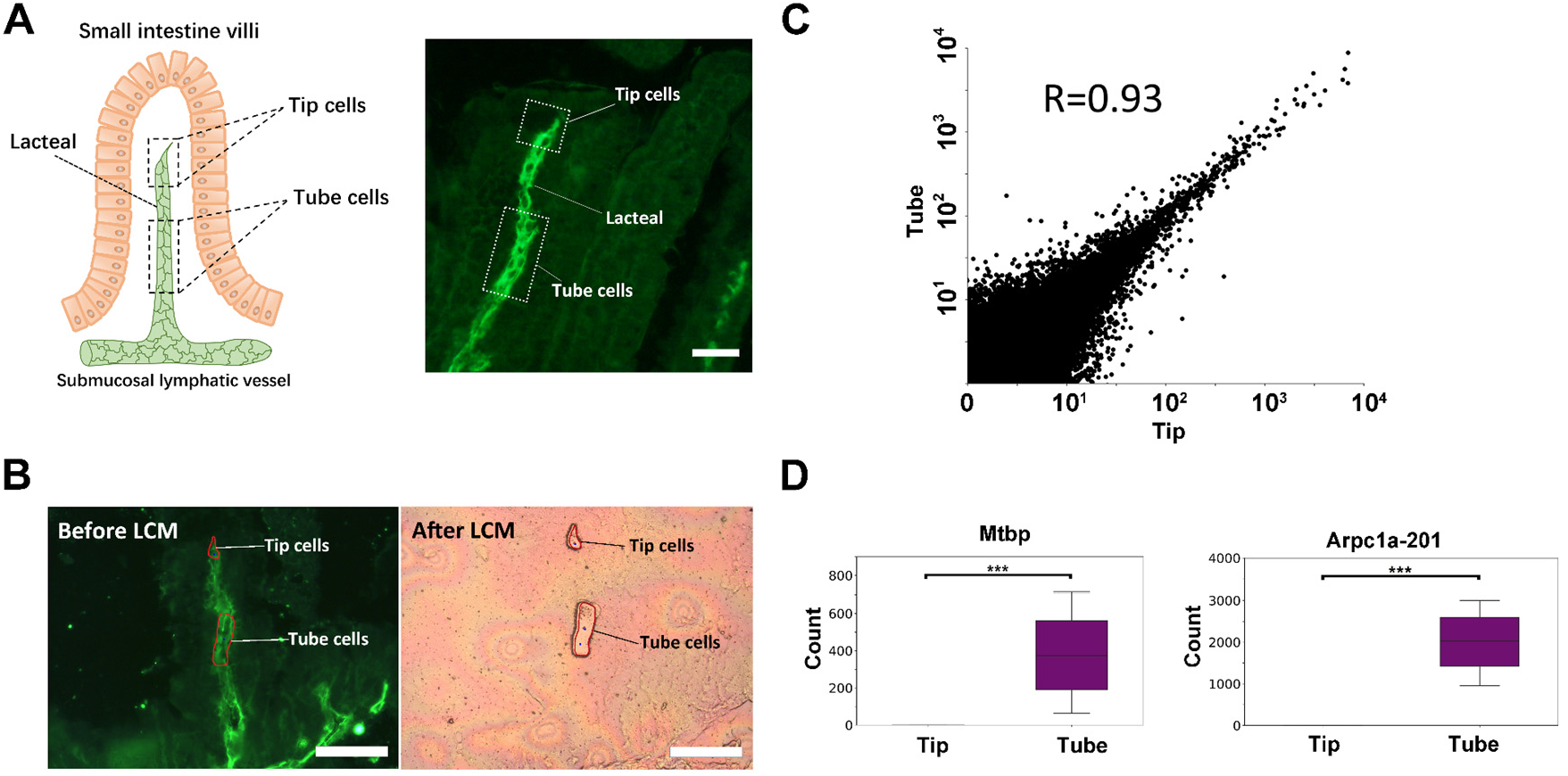
Transcriptome differences between the tip and the tube of the mouse small intestine lacteal. (**A**) Illustration of the small intestine lacteal structure (left) and a typical IF image of the lacteal (right), labelled with an anti-Lyve1 antibody. Scale bar: 50 μm. (**B**) Microdissection of the lacteal: tip cells and tube cells are collected separately with LCM. Each tip sample was pooled from up to 100 lacteals. Scale bar: 100 μm. (**C**) Scatter plot of the expression levels of the tip and the tube with all replicates combined. The highly consistent nature of the transcriptome is expected since these are all lymphatic endothelial cells. Although both the tip and the tube are considered to have the same phenotype, significant expression differences are still identified. Shown in (**D**) are two genes that are only expressed in the main body of the tube: *Mtbp* and the *Arpc1a-201* isoform (***, *p_adj_* < 10^-5^).

We then dissected a minute amount of material from the tip of the lacteals (~ 20 μm in length, equivalent to 1-2 cells). We pooled this material into three independent replicates, each with an estimated 150 cells. We obtained 50 – 58 million clean reads for each and the average overall mapping rate was 78% (Supplemental Table S2). As expected, the averaged Pearson coefficient between pairwise comparisons is 0.89 (Supplemental Fig. S9). Using FPKM ≥ 1, about 12,000 expressed genes were detected for each sample (Supplemental Table S2).

Combining all replicates, we explored the possibility of identifying differentially expressed genes or transcript isoforms using the program DESeq2. Despite the fact that the overall transcriptomes of the tip and the tube are highly similar (R = 0.93; Fig. 7C), which was expected since they are both lymphatic endothelial cells, DESeq2 was still able to identify several genes and transcript isoforms with a statistically significant difference in expression (Supplemental Fig. S10, Supplemental Table S5). Among these, the gene *Tnfsf15*, an established activator of lymphatic endothelial cell growth (Qin et al. 2015), was found to be only expressed in the tip of the lacteal (Supplemental Table S5), suggesting that it might be related to the proliferative characteristic of the lacteal cells. In addition, *Mtbp* and the protein-coding isoform of *Arpc1a* (201), both encoding for actin-filament severing proteins, were not expressed in the tip of the lacteal but robustly expressed in the tube cells (Fig. 7D, Supplemental Table S4-S5) (Agarwal et al. 2013; Abella et al. 2015). *Arpc1a* is highly conserved and its protein product is one of the components of the Arp2/3 complex that plays an essential role in generating branched actin filaments. The loss of *Arpc1a* was reported to result in long actin tails both in cells and *in vitro* (Abella et al. 2015), and thus, could be related to the presence of long filopodia-like protrusions at the lacteal tip. Although further work is needed to fully characterize the functional consequences of these differences in expression, these results provide a clear example of the power of spatially-resolved transcriptional analysis of a complex tissue. It should also be noted that only with transcript isoform analysis could the difference in *Arpc1a-201* expression in particular be detected with certainty for which high RNA quality is paramount.

### Extension of immuno-LCM-RNAseq to RNAlater-preserved tissues

While snap-freezing is the preservation method-of-choice for biological research, clinical tissues are often preserved with RNAlater owing to its convenience and potency to protect RNA during long-term storage at cryogenic temperatures (Florell et al. 2001; Mutter et al. 2004; Kasahara et al. 2006; Diaz et al. 2013). To enable full use of these resources, we further sought to extend our method to the tissues preserved with RNAlater (Fig. 8). However, one of the often encountered problems with RNAlater preserved tissues is a difficulty to section properly in a cryostat (Ellis et al. 2002; Legres et al. 2014). We found that this difficulty largely stemmed from the softness of the treated tissue at typical cryostat cutting temperatures (−20°C). We found that at ~ −24°C, the RNAlater solution alone freezes and stiffens significantly. Hence, by lowering the cutting temperature to ~ −28°C by externally introducing streaming liquid nitrogen gas across of the knife holder, we found that RNAlater preserved tissues could be routinely sectioned with sufficient robustness at the desired thickness without crumpling or sticking to the cutting blade. The second issue with RNAlater-preserved tissues is that its components are inhibitive to proper immunolabelling (Brown and Smith 2009), probably as a consequence of their interference with the antigen-antibody interaction. Therefore, IF labelling with these sections could only be performed after RNAlater was completely replaced with an RVC-containing solution. With these easily adaptable modifications, we were able to obtain high quality transcriptomes from IF-guided microdissection of tissues preserved by RNAlater. Similar to the snap-frozen tissues, we used both anti-Lyve1 and anti-cytokeratin (PanCK) antibodies to demonstrate the validity of the modified immuno-LCM-RNAseq protocol. As shown in Supplemental Fig. S11, in the presence of 10 mM RVC, high quality IF images were obtained with the mouse lacteal and stomach lymphatic vessels. With ~1,500 cytokeratin positive (panCK) cells from the crypt region of the mouse small intestine, we obtained ~15 million clean reads with an 82% overall mapping rate (Supplemental Table S2). About 14,000 expressed genes were detected at FPKM ≥ 1, similar to that with snap-frozen tissues (Supplemental Table S2). Furthermore, the transcriptomes from the two different preservation methods were also in high agreement with an average R value of 0.84 (Supplemental Fig. S12, Supplemental Table S2).

**Figure 8.**
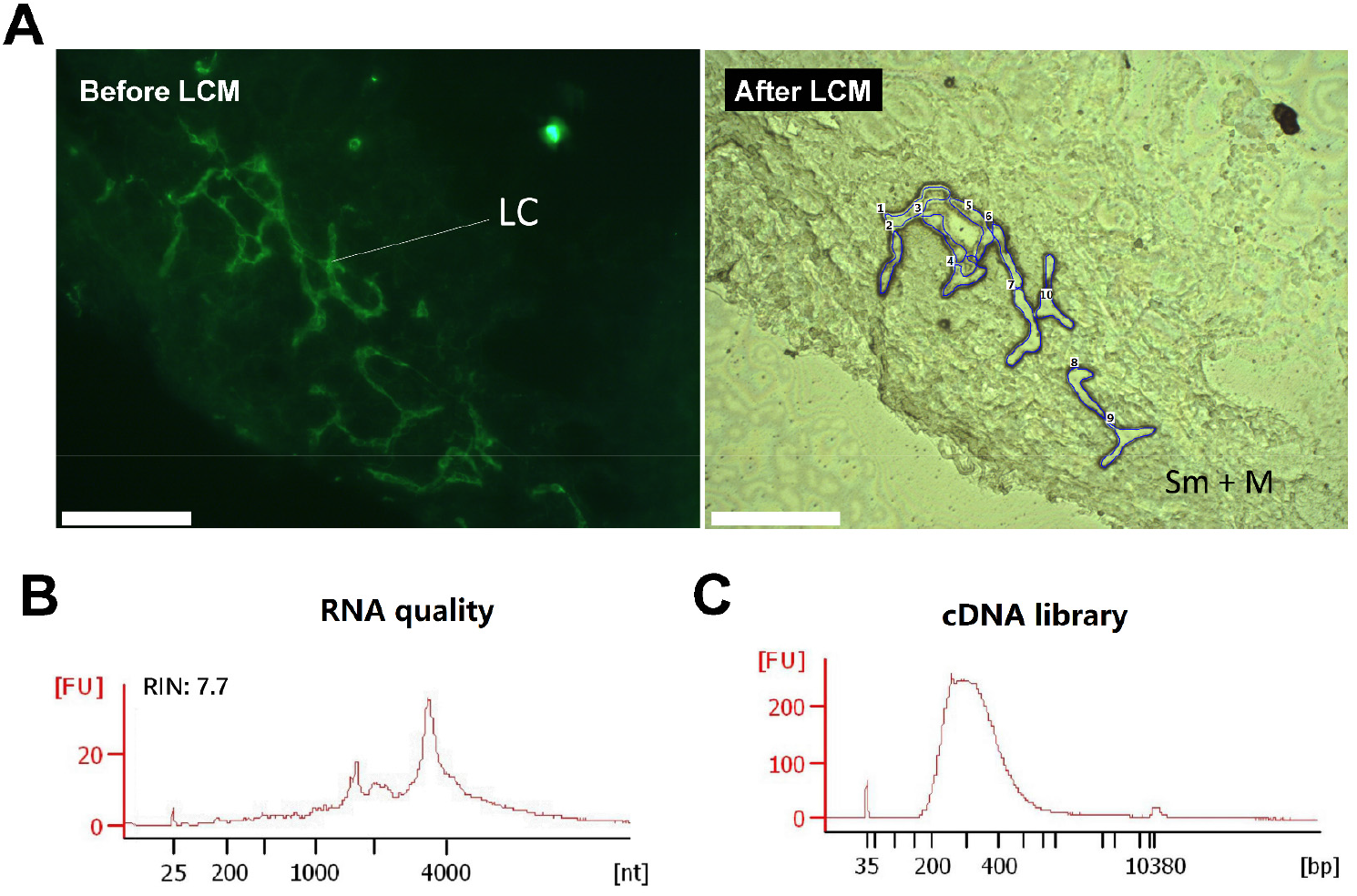
Immuno-LCM-RNAseq of the human jejunum lymphatic vessel from a clinical, RNAlater-preserved jejunum tissue. (**A**) Left: Immunofluorescence image of the lymphatic vessels in the human jejunum tissue section using an anti-Podoplanin antibody (with 10 mM RVC). Right: bright field image after LCM. LC: lymphatic cells; Sm: submucosal layer; M: muscle layer (**B**) RNA quality of the leftover materials after LCM. (**C**) The quality of the cDNA library prepared from this RNA. Scale bar: 200 μm.

As a further demonstration, we applied our approach to the analysis of an RNAlater preserved clinical sample, the human jejunum. Using the anti-Podoplanin antibody to identify lymphatic cells in these tissue sections, we micro-dissected ~100 Podoplanin-positive cells to profile their gene expression (Fig. 8). With 43 million clean reads, we were able to detect 15,000 genes expressed in these lymphatic endothelial cells (Supplemental Table S2 and S6). To lend further validity of this result, we compared our transcriptome to a known expression profile of the human primary dermal lymphatic endothelial cells which is based on bulk RNAseq (Breschi et al. 2020). The resulting Pearson correlation between these two samples is 0.89 (R=0.89), demonstrating the reliability of our method to characterize clinical samples.

## Discussion

Laser capture microdissection (LCM) (Emmert-Buck et al. 1996) offers a unique ability to precisely isolate targeted materials from either snap-frozen or RNAlater preserved bio-sections, even down to single cell levels. When combined with well-characterized phenotype markers, such as antibodies, it has long been known that LCM could provide detailed, spatially defined and cell-type guided transcriptome analysis with unparalleled precision. However, the key for the successful incorporation of this technology into the repertoire of spatial transcriptome methods is the protection of the RNA integrity during the lengthy process: the IF labelling often requires many hours to complete and the microdissection procedure could also take a long time to finish. In these processes, both endogenous and exogenous (such as airborne) RNases can quickly degrade the RNA exposed in the sectioned tissues. Although various schemes and protocols have been proposed to overcome this problem, the RNA quality issues remained unresolved until now. This is true even for formalin-fixed and paraffin-embedded (FFPE) tissue samples (Murray 2018; Foley et al. 2019). As demonstrated in this study, with the inclusion of a moderate amount of a small ribonuclease inhibitor, RVC, in most steps that must be performed under ambient conditions, RNA can be effectively protected with high quality RNA profiles acquired from as little as a few tens of dissected cells, mostly owing to their small size and fast diffusion in the dense tissue sections, which had not been explored before (de Heredia and Jansen 2004; Ohlson et al. 2005; Zielinski et al. 2006; Graindorge et al. 2019). It is as important that RVC has negligible effect on the actions of both primary and secondary antibodies and allows the immunolabelling to proceed according to the standard procedures. Our method also significantly outperforms results recently obtained using the Rapid protocol with bone marrow tissue (Baccin et al. 2020) in terms of number of genes as well as overall mapping rate, and rRNA, exonic, intronic, and intergenic mapping rates (Supplemental Fig. S13, Supplemental Table S2). Given the simplicity of our approach, we anticipate that immuno-LCM-RNAseq can be adopted with most existing LCM platforms, facilitating its use in refined analysis of the spatial transcriptomes of intricately structured, phenotype complex tissues in their native state. As exemplified in the spatially select analysis of the lacteal capillary in the mouse small intestine, critical differences at the transcript isoform level could be resolved with a sensitivity unravelled by any other approach at this spatial scale.

Although with the present approach, as few as ~60 cells were demonstrated to be sufficient to acquire high quality transcriptomes, there should still be room for further improvement. In our particular protocol, the RNA extraction process is associated with a certain amount of material loss, especially when the collected materials are extremely low. For example, for 100 cells, the total mRNA contained is probably less than half a nanogram. In this regard, adopting a single tube approach might further improve the lower limit of immuno-LCM-RNAseq substantially. Based on what has been documented in the literature (Soumillon et al. 2014; Tang et al. 2009), analysis at the single cell level might even be feasible. However, when pushed to such a limit, whether it is still possible to achieve full-length transcript and isoform analysis remains to be established.

It should also be noted that with either snap-frozen or RNAlater preservation, the tissues should still retain their native state if the protocols are performed properly. Therefore, immuno-LCM-RNAseq allows for the native transcriptome, *i.e.*, under in vivo conditions and environments, to be acquired. Since these will be largely identified with immuno-defined phenotypes, *i.e.*, through protein targets, these are not necessarily identical to those derived through the clustering of scRNAseq profiles, where it is the collective of the most variant transcripts, but not proteins, that are used for classification. How these two types of phenotypes or cell states are related should be an intriguing and also important question to pursue. Indeed, the potential usefulness of LCM to confirm or further expand observations obtained with scRNAseq has also been noted in several recent scRNAseq studies (Baccin et al. 2020; MacParland et al. 2018; Longo et al. 2021). Previous difficulties with LCM-RNAseq had prevented a robust comparative study of this aspect of phenotype examination. Now with this robust protocol, we expect many interesting discoveries to emerge in the future delineations of functional states of tissue defined cells. Along the same line, the native transcriptomes obtained using immuno-LCM-RNAseq can also serve as the basic cell characterizations in the interpretation of most recent coded bead-based methods for spatial transcriptomics. In comparison to the “standard” transcriptomes derived from scRNAseq profiles, the transcriptomes acquired with immuno-LCM-RNAseq are not only more comprehensive, but even more importantly, obtained under exactly the same conditions. In fact, potential artefacts owing to single cell preparation procedures notwithstanding (van den Brink et al. 2017), even the best separated clusters in a single cell dataset cannot provide a detailed profile with isoforms quantitatively resolved as in the immuno-LCM-RNAseq transcriptomes.

In summary, in overcoming the long-standing problem to preserve RNA quality, we have established a novel, powerful method of spatial transcriptomics based on immuno-guided LCM that is complementary to existing approaches, as well as unique with its own significant advantages. Rare cell types in particular or cells whose functioning is exquisitely sensitive to their positioning or environment within the tissue can now be interrogated to obtain an understanding of their complete transcriptome, including transcript isoforms. There is no doubt that it is only with such a systems-wide characterization of the cells within the native tissue can their functioning be fully understood, which is ultimately essential for an understanding of the functioning of the tissues as a whole, whether healthy or diseased.

## Methods

### Preparation of tissues

6-8 weeks old C57BL/6 mice (Jie Si Jie Laboratory Animals, Shanghai, China) were used in these experiments. All mice were euthanized by cervical dislocation. The stomach, small intestine, liver, kidney, colon, spleen and testis were freshly dissected and cleaned in ice-cold RNase-free phosphate-buffered saline (PBS) solution. After absorption of excess liquid, tissues were placed in 2.5 ml capped cryogenic vials individually, and sealed with Parafilm (M2 Scientifics, cat.no. HS234526BC, Holland, Michigan, USA). Tissues were then snap-frozen in liquid nitrogen for 30 min and stored at −80°C.

For samples preserved in RNAlater (Invitrogen, cat.no. 7021, Carlsbad, California, USA) which protects RNA from degradation, tissues were first cut into pieces smaller than 0.5×0.5×0.5 cm^3^ and washed in ice-cold RNAlater solution, then left in the RNAlater solution at a RNAlater:sample volume ratio ≥ 10:1 at 4°C for 4 to 8 hrs. Excess RNAlater solution was then removed and the treated tissue was stored at −80°C. All experiments with animals were performed in compliance with the guidelines of the Institutional Animal Care and Use Committee of the Shanghai Jiao Tong University. The clinical human jejunum tissue was collected from a gastrectomy patient (Renji Hospital, Shanghai, China). The tissue was cut into small pieces and immediately placed into the RNAlater solution on ice. Tissue samples were stored at −80°C until use. Approvals were obtained from the Research Ethics Committee at Renji Hospital, Shanghai, China.

### Sectioning of tissues with the cryostat

For snap-frozen tissues, the cryostat (Leica, cat.no. CM3050S, Buffalo Grove, Illinois, USA) was first defrosted and spray-cleaned with RNaseZap (Ambion, cat. no. AM9780, Austin, Texas, USA) and pure ethanol (Sigma-Aldrich, cat.no. E7023, St. Louis, Missouri, USA). The cryochamber temperature (CT) was set at −22°C and the specimen temperature (OT) was set at −20°C as commonly recommended (Dey 2018). The Optimal Cutting Temperature compound (OCT) (Agar Scientific, cat. no. AGR1180, Stansted, UK) was placed on ice for at least 30 min before use. A PET (polyethylene terephthalate)-membrane covered slide (Carl Zeiss, cat.no. 415190-9051-000, Jena, Germany) was cleaned with RNaseZap followed by UV irradiation for 30 min prior to mounting the sections. Most tissues were embedded in OCT and cut into 12 μm-thick sections, and mounted on PET slides. Sections on PET slides were dried for 2 to 3 min in the cryostat before fixation (for IF). For the mouse small intestine, it was critical to properly orient the tissue in the OCT in order to obtain desired sections. To this end, a frozen OCT block was first cut to obtain a flat surface, taking note of the cutting direction. The intestine tissue was then re-embedded on this flat surface with OCT, cut into 12 μm-thick serial sections and mounted on a PET slide.

For RNAlater-preserved tissues, the CT and OT were kept below −28°C. This was achieved by flowing a stream of liquid nitrogen gas across the cryostat knife holder. The frozen RNAlater-preserved tissue often contained a thin layer of solidified RNAlater material on the surface, which prevented direct contact between the tissue and OCT, and often led to difficulties in proper sectioning. To remove this solidified RNAlater layer, we partially immersed the frozen RNAlater-preserved tissue in fresh ice-cold OCT, and after OCT solidified, removed the sample from the frozen OCT, which left a large portion of the RNAlater layer attached to the frozen OCT. This was performed repeatedly until the residual RNAlater layer was completely removed from the tissue. The tissue was then fully embedded within ice-cold OCT, cut into 12 μm-thick serial sections and mounted on a PET slide. The sections were dried for 2 to 3 min in the cryostat before fixation (for IF).

### Immunostaining with the Rapid protocol and the high-salt protocol

All immunostaining procedures were performed in a RNA-specific biological safety cabinet which was pre-cleaned by RNaseZap. However, antibody labelling was carried out at 4°C in a refrigerated chamber.

For the Rapid protocol, we followed a previously established procedure (Nichterwitz et al. 2016). The air-dried (in the cryostat) PET slide with the sections was fixed for 5 min in ice-cold acetone (Sigma-Aldrich, cat.no. 179124, St. Louis, Missouri, USA) followed by 3 quick washes (1 min) with RNase-free ice-cold PBS solution. The slide was then incubated with rabbit anti-mouse Lyve1 primary antibody (1:25, AngioBio cat.no. 11-034, San Diego, California, USA) in cold PBS with 0.25% Triton X-100 (Sigma Aldrich, cat.no. 93426, St. Louis, Missouri, USA) for 5 min at 4°C. After 3 quick washes with ice-cold PBS (1 min), the slide was incubated with secondary antibody (1:25; Alexa Fluor 488 conjugated goat-anti-rabbit; Invitrogen, cat.no. A11034, Carlsbad, California, USA) in ice-cold PBS with 0.25% Triton X-100 for 5 min at 4°C and then washed 3 times in ice-cold PBS (1 min).

For the high-salt protocol, we closely followed the original protocol presented in the ref (Brown and Smith 2009). In short, the sections were fixed in 70% ethanol for 5 min, then followed with a rapid PBS wash. Sections were incubated with rabbit anti-mouse Lyve1 antibody (1:100, AngioBio cat.no. 11-034, San Diego, California, USA) with 2 M NaCl in PBS overnight at 4°C. Unbound primary antibody was removed by 3 quick washes with 2 M NaCl in PBS for 5 min. Sections were then incubated with Alexa Fluor 488 conjugated goat-anti-rabbit (1:100, Invitrogen, cat.no. A11034, Carlsbad, California, USA) in 2 M NaCl PBS in the dark for 1 hour at 4°C. Slides were then washed 3 times in 2 M NaCl PBS for 5 min. All of these solutions were ice-cold. We note that the original procedure used overnight incubation with the antibodies. Owing to the serious structural damage on the small intestine sections under this condition (see Supplemental Fig. S1), we also examined a shorter incubation procedure similar to that described above except with a ~3.5 hrs primary antibody labelling (~5 hrs. total incubation time). To reduce cross-reactivity, we also examined a procedure that included a blocking step, prior to incubation with the primary antibody, using a mixture of equal volume of ready-to-use protein block serum-free solution (Agilent Dako, cat.no. X0909, Santa Clara, California, USA) and 4 M NaCl in PBS at 4°C for 15 min followed by 3 quick washes with 2 M NaCl in PBS. In those experiments with this blocking step, we also diluted the antibodies in the 2 M NaCl solution and a 1:4 dilution of the ready-to-use protein block serum-free solution.

### RVC-based immunofluorescence staining procedure

The sections on PET slides were first fixed with cold acetone in the cryostat: 30 s for snap-frozen tissues and 5 s for RNAlater-preserved tissues. The fixed sections were then dried in the cryostat for 5 min, and washed 3 times with ice-cold 10 mM Ribonucleoside Vanadyl Complex (RVC) (New England BioLabs, cat.no. S1402S, Ipswich, Mass, USA) in buffer A (10 mM NaCl, 3 mM MgCl_2_, 20 mM Tris•HCl, pH 7.4) in a RNA-specific biological safety cabinet. We refer to this 10 mM RVC solution as the RVC solution unless otherwise indicated. The sections were pre-blocked with an equal volume mixture of 20 mM RVC in buffer A and the ready-to-use protein block serum-free solution at 4°C for 15 min, followed by 3 times wash with the RVC solution. The slides were then incubated for 3.5 hrs with the primary antibody: either anti-mouse Pan Cytokeratin antibody (PanCK) (1:100, Santa Cruz Biotechnology, cat.no. sc-8018, Dallas, Texas, USA), anti-mouse Lyve1 antibody (1:100, AngioBio cat.no. 11-034, San Diego, California, USA) or anti-human Podoplanin antibody (1:100, ReliaTech, cat.no. 101-M41, Wolfenbuettel, Germany). After washing 3 times with the RVC solution, the slides were incubated with the secondary antibody, either Alexa Fluor 488 conjugated goat-anti-rabbit or donkey anti-mouse (1:100, Invitrogen, cat.no. A11034, A32766 Carlsbad, California, USA) or Fluor 568 conjugated donkey anti-mouse (1:100, Invitrogen, cat.no. A10037), in the dark for 1 hr at 4°C. All antibodies were pre-diluted in the RVC solution and a 1:4 dilution of the ready-to-use protein block serum-free solution before use. After secondary antibody incubation, the slides were washed 10 times with the RVC solution and temporarily stored in a light-tight box until laser microdissection. For validation of the lymphatic vessel location in the small intestine or stomach tissue sections, we further stained with Hoechst (1:1000 in RVC solution, Invitrogen, cat.no. H3569, Carlsbad, California, USA) in the dark at 4°C for 10 min, then washed 10 times with the ice-cold RVC solution. For maximum RNA protection, the RVC solutions must be freshly prepared before use: long term storage, even at 4°C, can severely reduce its effectiveness.

### Laser capture microdissection of immunofluorescence identified cells

Laser capture microdissection was performed with the Zeiss PALM MicroBeam LCM system (Zeiss Microimaging, Munich, Germany) housed in a Plexiglas housing. After the desired cells were manually selected on the control screen based on their fluorescence signal, excess solution was removed with a pipette from the slide and the sections were allowed to dry fully on the PALM stage under controlled humidity (humidity < 45%). The dissected materials with the UV laser were ejected into 0.2 ml adhesive cap tubes (Carl Zeiss, cat.no. 415190-9191-000, Jena, Germany). The tubes were quickly removed and taken to a Biological Safety Cabinet (Thermo Fisher, 1300 Series A2) that was pre-cleaned with RNaseZap. 30 μl of GITC lysis buffer (Invitrogen, cat.no. 15577-018, Carlsbad, California, USA) was added to the cap with gentle pipetting. The tubes were then sealed with Parafilm and vortexed several times, followed by incubation at 42°C for 30 min to improve RNA extraction. The tubes were then centrifuged at 20,800g for 10 min at room temperature and stored at −80°C for later use.

### Assessment of RNA quality

For all of the following, RNA was extracted using the RNeasy Micro Kit (QIAGEN, cat.no. 74004, Hilden, Germany) and examined with the 2100 Bioanalyzer (RNA6000 Pico Kit, Agilent, cat.no. 5067-1513, Santa Clara, California, USA). The RNA quality of the tissues was initially evaluated by extracting the total RNA from a few pieces of the sections of either the snap-frozen or RNAlater preserved tissues. Only those tissues with an RNA integrity number (RIN) greater than 9.0 were used. The RNA quality was also evaluated after microdissection by examining the leftover materials from the same section to make sure that there was no serious degradation during the procedure. Only those with RIN > 7.0 were considered of sufficient quality for further analysis. As documented in literature, an RIN > 6.5 is generally considered sufficient for transcriptomic analyses as lower RIN samples often result in the loss of library complexity (Romero et al. 2014). These leftover materials were collected by LCM and placed into 350 μl of RLT lysis buffer (RNeasy, Qiagen, including 1%β-Mercaptoethanol (β-Me)) followed by RNA extraction and examination with the 2100 Bioanalyzer.

### Preparation of the cDNA library

The samples stored in the GITC lysis buffer were thawed on ice, pooled, and then additional GITC lysis buffer was added to bring the total volume to 200 μl. RNA/DNA was precipitated by incubation at −80°C for 2 hrs with 600 μl cold ethanol (Sigma-Aldrich, cat.no. E7023, St. Louis, Missouri, USA), 20 μl 3 M NaAc (Amresco, cat.no. 97062-812, Soren, Ohio, USA) and 1 μl Glycogen (Invitrogen, cat.no. R0551, Carlsbad, California, USA). The samples were then centrifuged for 30 min at 4°C and the precipitate was washed 3 times with 75% cold EtOH, followed by dissolution in 10 μl RNase-free water (including 2U/μl SUPERase• In™ RNase Inhibitor). The DNA was digested with HL-dsDNase in DNase buffer (NuGEN, cat.no. 0354, San Carlos, California, USA) at 39°C for 15 min. The cDNA library was constructed using the Ovation SoLo RNAseq Kit (NuGEN, cat.no. 0354, 0352, San Carlos, California, USA), according to the manufacturer’s instructions. The number of optimal PCR cycles was determined by qPCR following the manufacturer’s recommendations. The cDNA library quality was evaluated using the 2100 Bioanalyzer (DNA high sensitivity kit, Agilent, cat. no. 5067-4626, Santa Clara, California, USA).

### RNAseq and data analysis

The cDNA library was sequenced on the Illumina high-throughput sequencing platform with the 2×150 bp pair-end mode. Raw reads were first submitted to Cutadapt-1.16 (Martin 2011) (with parameters of --u 5 --max-n 0 --minimum-length 100) to remove the sequencing adapters. The first 5 bases of each read were removed according to the library construction protocol of the Ovation SoLo RNAseq Kit. Trimmomatic-0.35 (Bolger et al. 2014) (with parameters of PE SLIDINGWINDOW:3:10 LEADING:10 TRAILING:10 MINLEN:100) was employed to remove low quality reads. SortMeRNA-v2.1b (Kopylova et al. 2012) was used to remove rRNA reads in the pair-end mode with default parameters. The cleaned reads were manually inspected by the Q30 profile of FastQC-v0.11.5 (Andrews 2010) to ensure sufficient data quality for further analysis. For the mouse data, the cleaned reads were mapped to the mouse GRCm38 (mm10) genome assembly with hisat2-2.0.5 (Kim et al. 2015; Pertea et al. 2016) in a strand-specific manner (with parameters of --rna-strandness FR). For the human data, the cleaned reads were mapped to the human GRCh38 (hg38) genome assembly with hisat2-2.0.5 (Kim et al. 2015; Pertea et al. 2016) also in a strand-specific manner (with parameters of --rna-strandness FR).

To evaluate the reproducibility of the replicates, the mouse genome was partitioned into 1 kb bins and the number of clean reads in each bin was counted with bedtools (v2.27). The Pearson correlation coefficient was then calculated pairwise between the samples. Transcript and gene level expression was estimated with StringTie-1.3.3 (Pertea et al. 2016; Pertea et al. 2015) (with parameters of -e -b) based on the Ensembl gene model (Mus_musculus.GRCm38.94.gtf and Homo_sapiens.GRCh38.94.gtf). Uniquely mapped clean read counts were normalized into FPKM (fragments per kb per million) to quantify gene and transcript expression.

The correlation between mouse small intestine lacteal tip and tube cells was calculated by using the average gene expression levels of the combined data of all replicates for each. Differential expression was examined between the tube samples (4 replicates) and the tip samples (3 replicates) using the DESeq2 package (1.28.1) (Love et al. 2014) with default settings. A subprogram prepDE.py of StringTie was employed to derive hypothetical read counts for each gene or transcript and the derived reads count matrix served as the input file of DESeq2 to conduct differential analysis at both the gene and transcript isoform level. Genes or transcripts with a log_2_FoldChange > 2 or log_2_FoldChange < −2 and p_adj_ < 0.01 were considered differentially expressed. Hierarchical analysis was performed by measuring the average Euclidean distance between different clusters. Gene body coverage plot was generated by RSeQC (v3.0.1) (Wang et al. 2012).

For the analysis of the data from the recent LCM RNAseq study of bone marrow (Baccin et al. 2020), we randomly selected nine samples with ~20 million reads (as examples with a comparable read depth as our small intestine epithelial samples) and four samples with ~30 million reads (as examples with ~50% greater read depth as our samples). Each sample was analysed using the same pipeline as our data (see above) except that parameters within sortMeRNA-v2.1b (Bolger et al. 2014) and hisat2-2.0.5 (Andrews 2010; Kim et al. 2015) were adjusted for processing of these 75 bp-single end datasets rather than our paired-end datasets.

## Data Access

All raw and processed sequencing data generated in this study have been submitted to the NCBI Gene Expression Omnibus (GEO; https://www.ncbi.nlm.nih.gov/geo/) under accession number GSE158396.

The RNAseq data generated in this study have been submitted to the NCBI BioProject database (https://www.ncbi.nlm.nih.gov/bioproject/) under accession number PRJNA658865.

List of genes and transcripts derived from differential expression analysis can be found in Supplemental Table S5. Gene and transcript level expression data of human jejunum lymphatic vessel cells can be found in Supplemental Table S6.

## Author’s contributions

Z.S. conceived the basic concept and X.Z., Y.G., D.M.C., Z.S. designed the experiments. X.Z performed the experiments. C.S.H. and H.L. performed the data analysis. C.H. provided the human jejunum samples. Y.W. and M.H. collected mouse colon and testis tissues. X.L. participated in imaging experiments. *J.W.* participated in data analysis and manuscript preparations. X.Z., C.S.H., Y.G., D.M.C. and Z.S. drafted the manuscript. All authors read and approved the final version of the manuscript. This work was supported by National Key R&D Program of China (Grant No. 2018YFC1003500), National Natural Science Foundation of China (Grant Nos. 81972909, 31670722, 31971151, 81627801), the Bio-ID Center at SJTU and the K. C. Wong Education Foundation (H. K.).

