## Supplemental Figure 1-12 and Supplemental Table 1-4 for "Robust Acquisition of High Resolution Spatial Transcriptomes from Preserved Tissues with Immunofluorescence Based Laser Capture Microdissection"

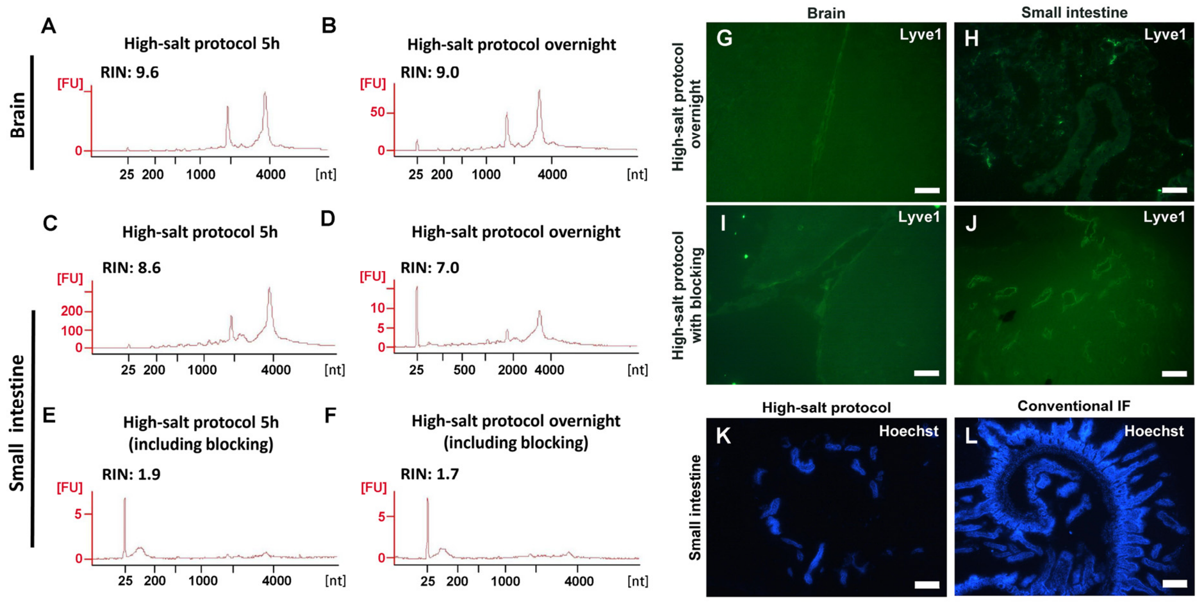


**Figure S1: Effectiveness of the high-salt protocol to preserve RNA and produce high quality IF images of frozen tissue sections.** (**A-D**) RNA quality of the mouse brain (**A, B**) and small intestine (**C, D**) frozen sections after a 5 hours (**A, C**) or overnight (**B, D**) high-salt (2 M NaCl) protocol. The original protocol described an overnight IF staining. This shorter, 5 hours-incubation was also examined here owing to the poor quality of the IF images on small intestine sections with the overnight incubation (**H**). (**E, F**) RNA quality of mouse small intestine frozen sections after the 5 hours (**E**) or overnight (**F**) high-salt protocol, including a blocking step in the staining procedure, showing degraded RNA. The original protocol did not include a blocking step, which might have contributed to the poor quality of IF images, and so we also examined this high-salt protocol with a blocking step. (**G, H**) IF images of the labelled lymphatic vessel cells in mouse brain (**G**) and small intestine (**H**) frozen sections following the overnight high-salt protocol, showing a lower signal-to-noise of the labeled cells than in the control (**Figure 1G-H**). (**I, J**) High-salt protocol labelled lymphatic vessel cells in the mouse brain (**I**) and small intestine (**J**) frozen sections, including a blocking step (~ 5 hours for the total procedure). (**K, L**) Images of the whole mouse small intestine tissue section after ~ 5 hours high-salt (**K**) or conventional IF (**L**), labelled by Hoechst. In the small intestine tissue sections, the high-salt protocol damaged the tissue section structure thus preventing further analysis. Scale bar: 50 µm (**G-J**), 200 µm (**K, L**).

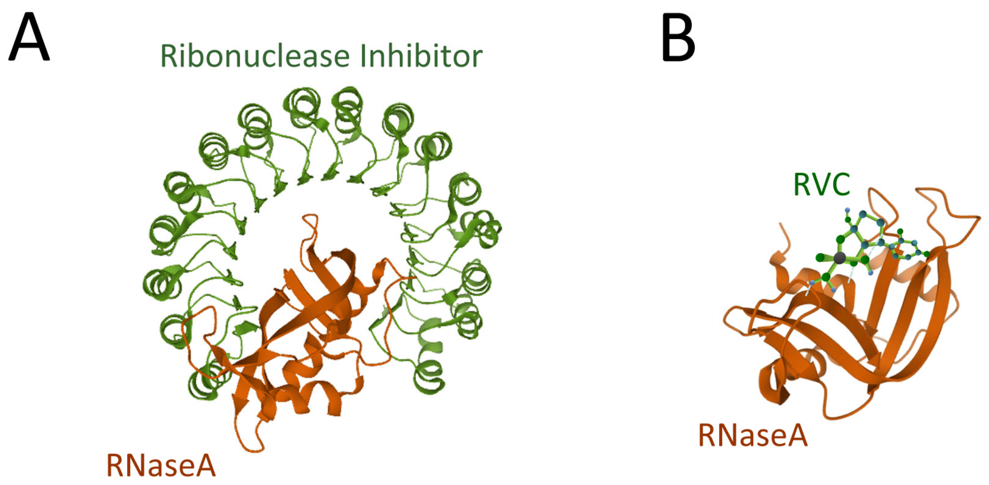


**Figure S2: Structural models depicting the interaction of RNaseA with inhibitors.** (**A**) The RNase inhibitor bound to RNaseA (pdb: 3TSR). (**B**) Model of RVC interacting within the active site of RNaseA to give a sense of the relative differences between the RNase inhibitor and RVC. This model was generated by manually positioning a typical component of RVC (the uridine vanadyl complex) within the active site of RNaseA.

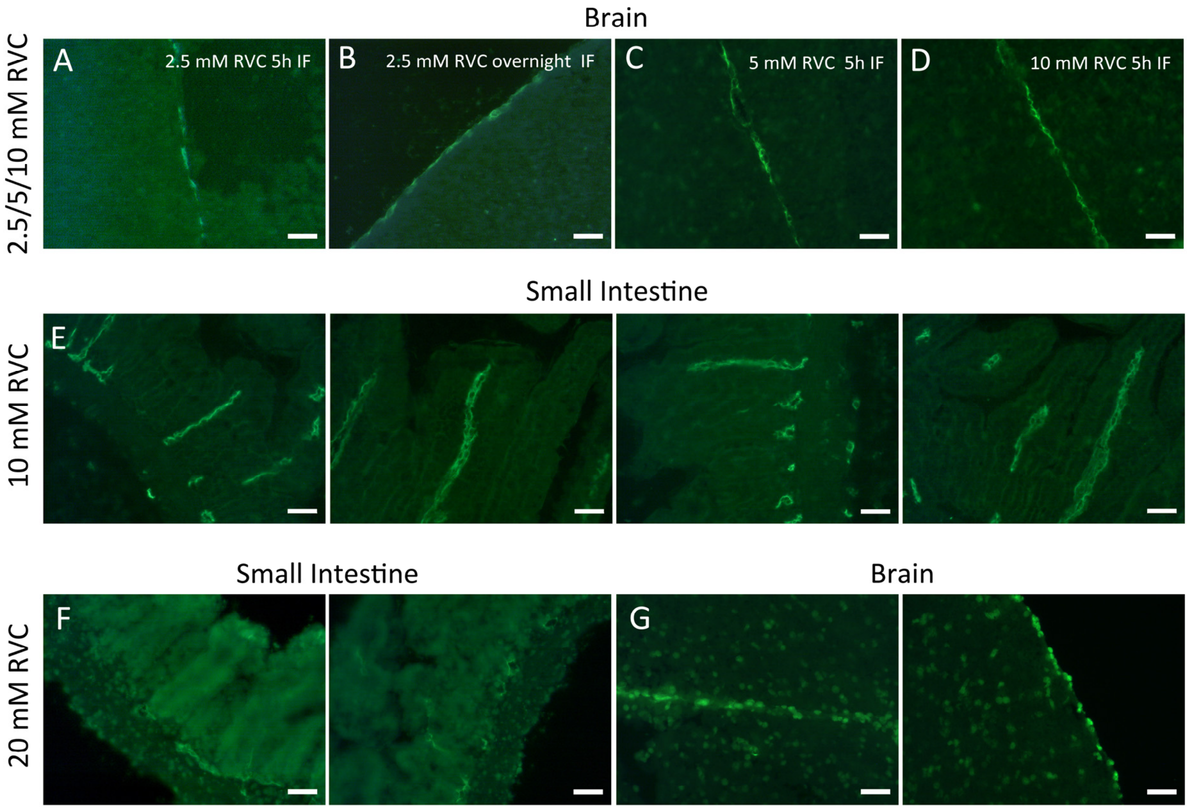


**Figure S3: Quality of IF images following IF in the presence of RVC with different frozen tissue sections.** (**A, B**) Images of mouse brain sections following a 5 hours (**A**) or overnight (**B**) IF staining in the presence of 2.5 mM RVC. Note that the image quality of 5 hours IF staining is as good as overnight incubation. (**C, D**) Images of mouse brain sections using 5 mM (**C**) or 10 mM (**D**) RVC during IF, showing that there is no negative effect of RVC on image quality (compare with **Figure 1H**). (**E**) Several examples of IF images of mouse small intestine sections following the RVC-based IF procedure (in 10 mM RVC). (**F, G**) Typical IF images obtained of small intestine (**F**) or brain (**G**) sections in the presence of 20 mM RVC following the RVC-based IF procedure. Note that there is higher background and less distinct labeling in these images compared to the 10 mM RVC-based IF procedure shown in (**E**). All samples were stained with anti-Lyve1 antibody. Scale bar: 50 µm.

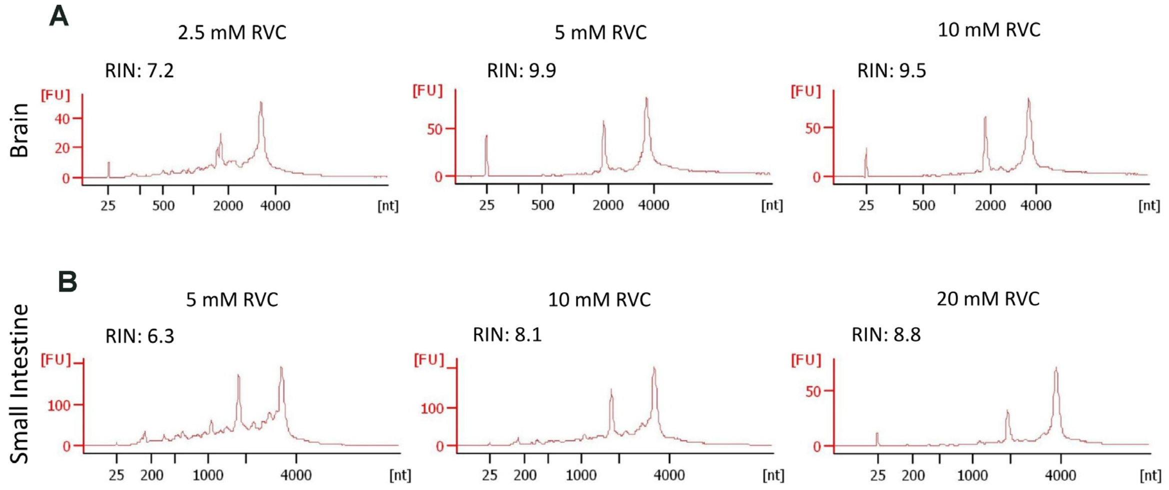


**Figure S4: Effectiveness of RVC to maintain RNA quality during IF staining of mouse brain and small intestine frozen tissue sections.** (**A, B**) The RNA quality of mouse brain (**A**) and small intestine frozen sections (**B**) in the presence of the indicated concentrations of RVC following IF (see Online Method). For brain tissues with low endogenous RNase, 5 mM RVC is sufficient to protect the RNA during the ~5 hours immunolabelling procedure, while for small intestine sections that have moderate levels of RNase, 10 mM RVC is needed to sufficiently preserve the RNA.

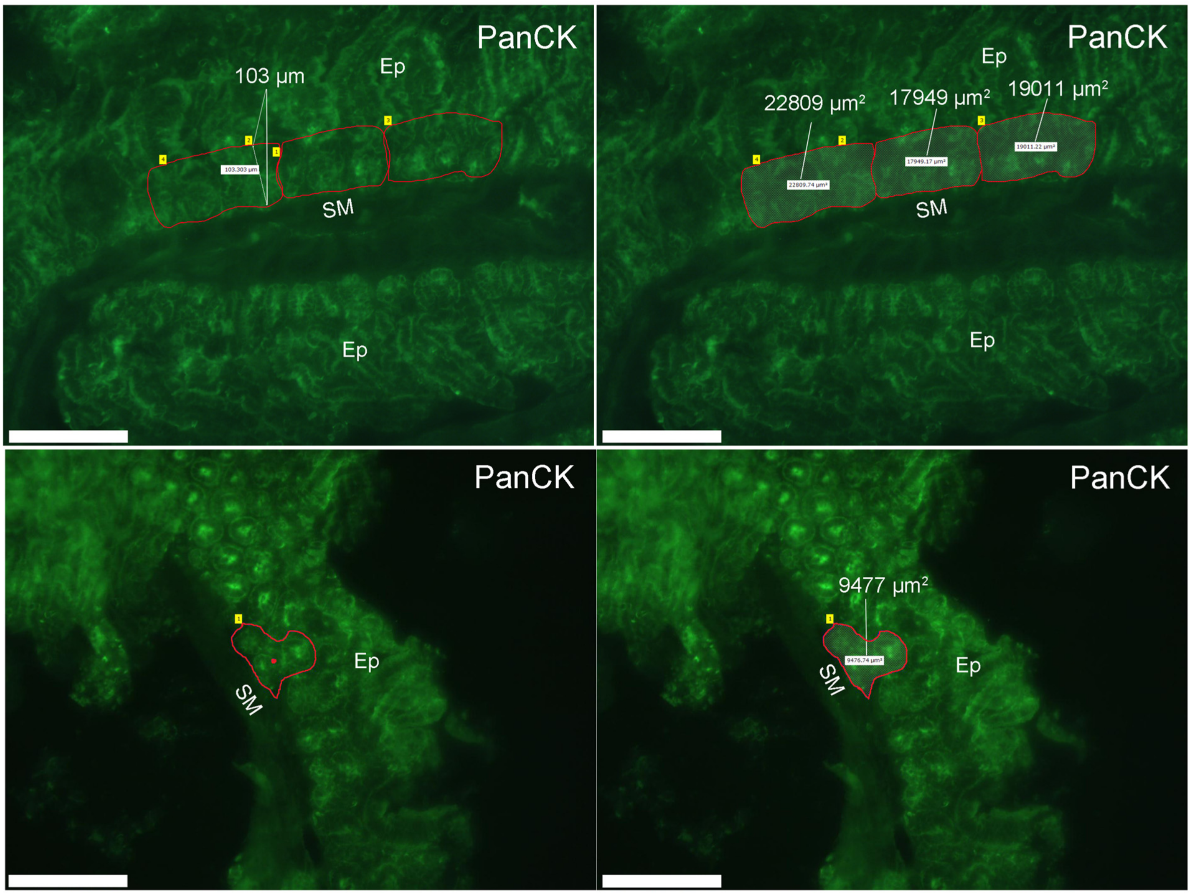


**Figure S5: IF images of mouse small intestine epithelial cells used to calculate the cell numbers of laser-dissected samples.** Examples of the IF images of PanCK-labelled epithelial cells (Ep) that were used to determine number of collected cells (see Supplementary Table 1). Ep cells only within ~100 µm of the submucosal layer (SM) were collected. Scale bar: 200 µm.

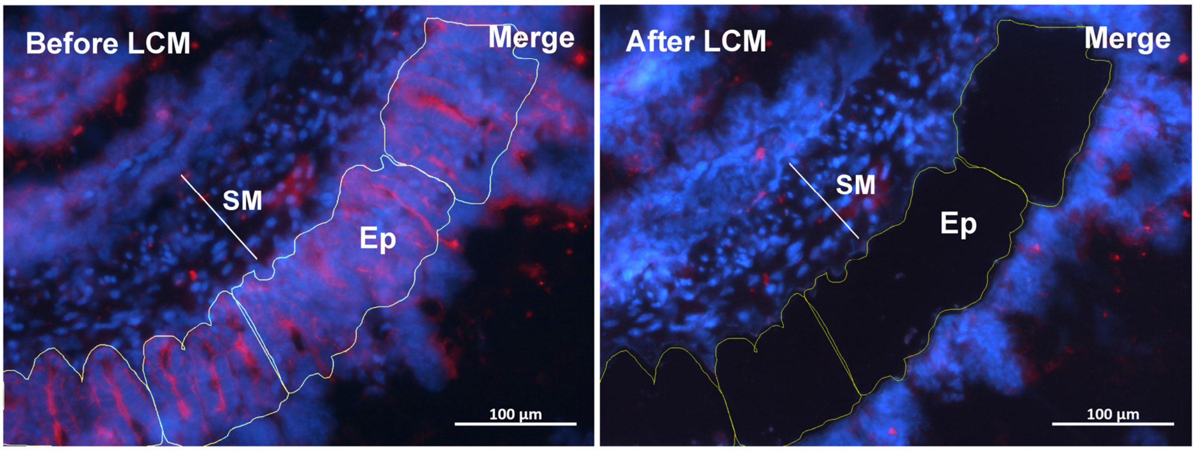


**Figure S6: Laser microdissection of the cytokeratin positive epithelial cells (Ep) in the mouse small intestine tissue section.** This image is of the same region as shown in **Figure 5** but here showing the nuclei of all cells (Red: PanCK, Blue: Hoechst). SM, submucosal layer; Ep, epithelial cells.

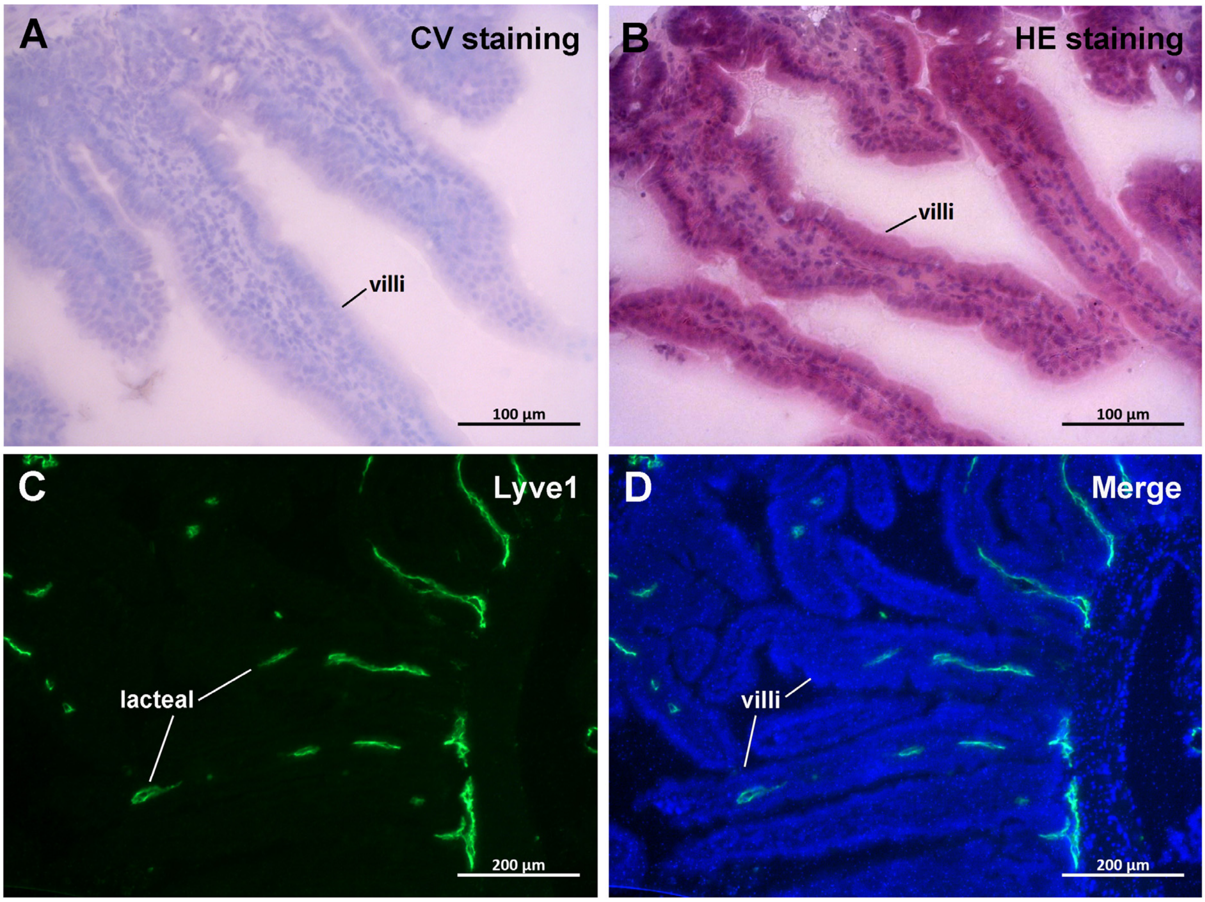


**Figure S7: Histological staining and immunolabelling images of mouse small intestine villi and lacteal.** (**A, B**) Cresyl violet and H&E staining of mouse small intestine sections. It is not possible to resolve the lacteal in this image. (**C, D**) Identification of the lacteal in the villi by immunostaining with the anti-Lyve1 antibody (Green: Lyve1, Blue: Hoechst).

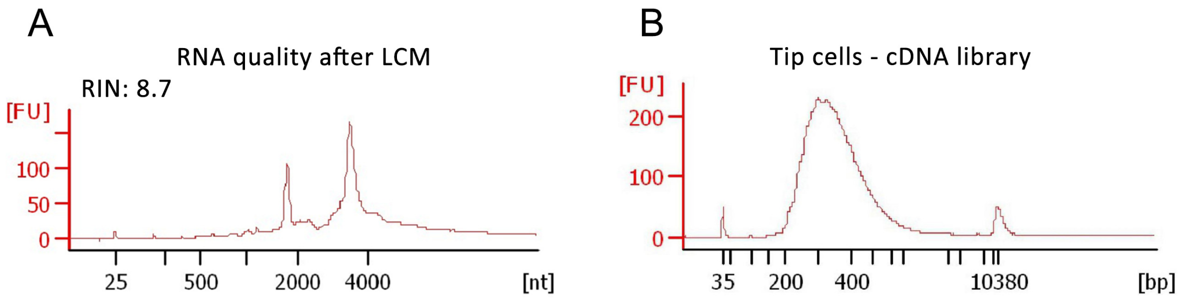


**Figure S8: RNA and cDNA library quality check following the immuno-LCM steps during the collection of the mouse lacteal tip cells.** (**A**) The RNA is well-preserved during this process. (**B**) The quality of the cDNA library generated from this RNA data is also high, as evidenced by the single distribution of sizes around 300 bp as expected.

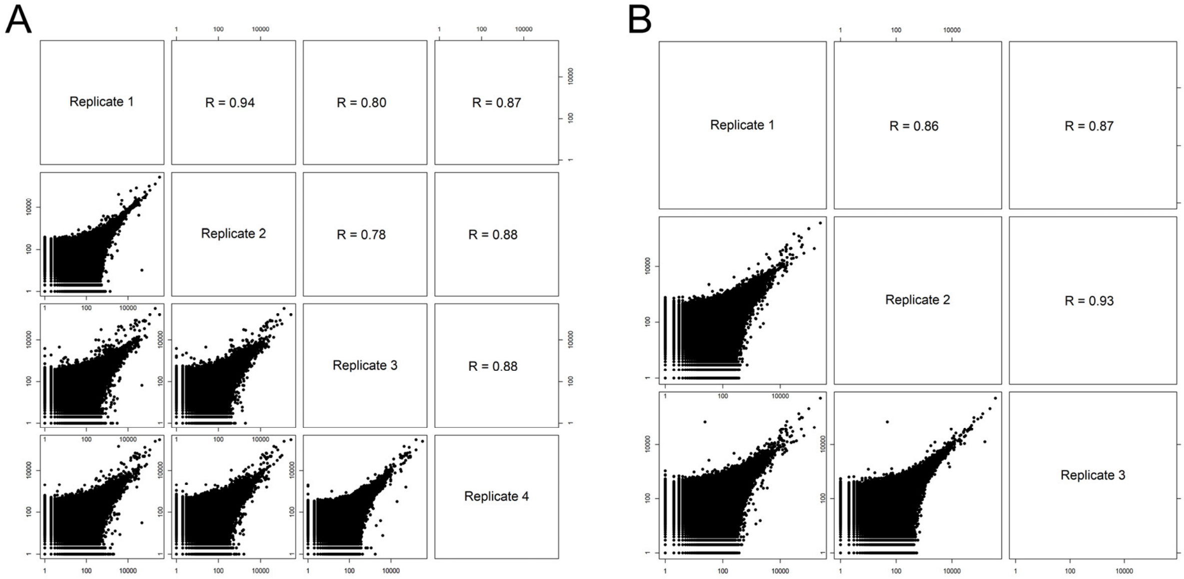


**Figure S9: Reproducibility of the RNAseq data among lacteal tip or tube replicates samples.** (**A, B**) Log-log scatter plots and Pearson correlation coefficients among the replicates of the lymphatic lacteal tube cell samples (**A**) and tip cell samples (**B**) from mouse small intestine frozen sections.

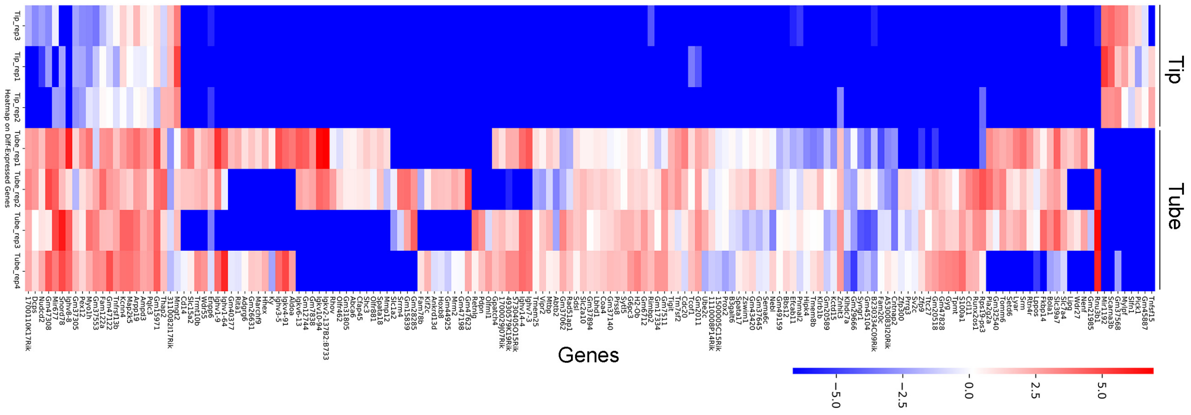


**Figure S10: Differentially expressed genes between tip and tube samples.** The 167 differentially expressed genes ($\log_2$(foldchange) > 2 or < -2, $p_{adj}$ < 0.01) between the tip and the tube cells from the mouse small intestine lacteals.

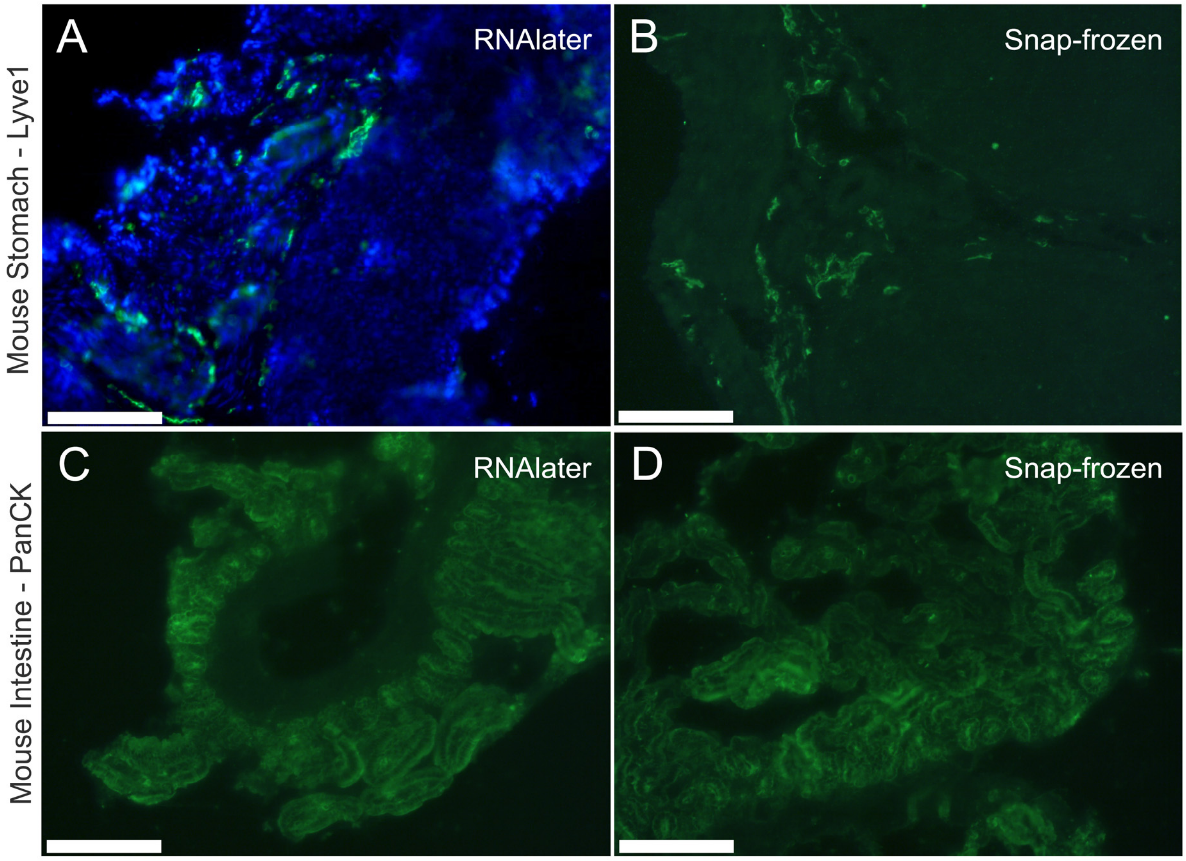


**Figure S11: IF image quality following the RVC-based IF with RNAlater-preserved or snap-frozen tissue sections.** (**A, B**) Lymphatic vessels labeled with anti-Lyve1 antibody from RNAlater-preserved (**A**) and snap-frozen (**B**) mouse stomach sections. (**C, D**) Epithelial cells labeled with anti-cytokeratin antibody (PanCK) with RNAlater-preserved (**C**) and snap-frozen (**D**) mouse small intestine sections. Scale bar: 200 μm.

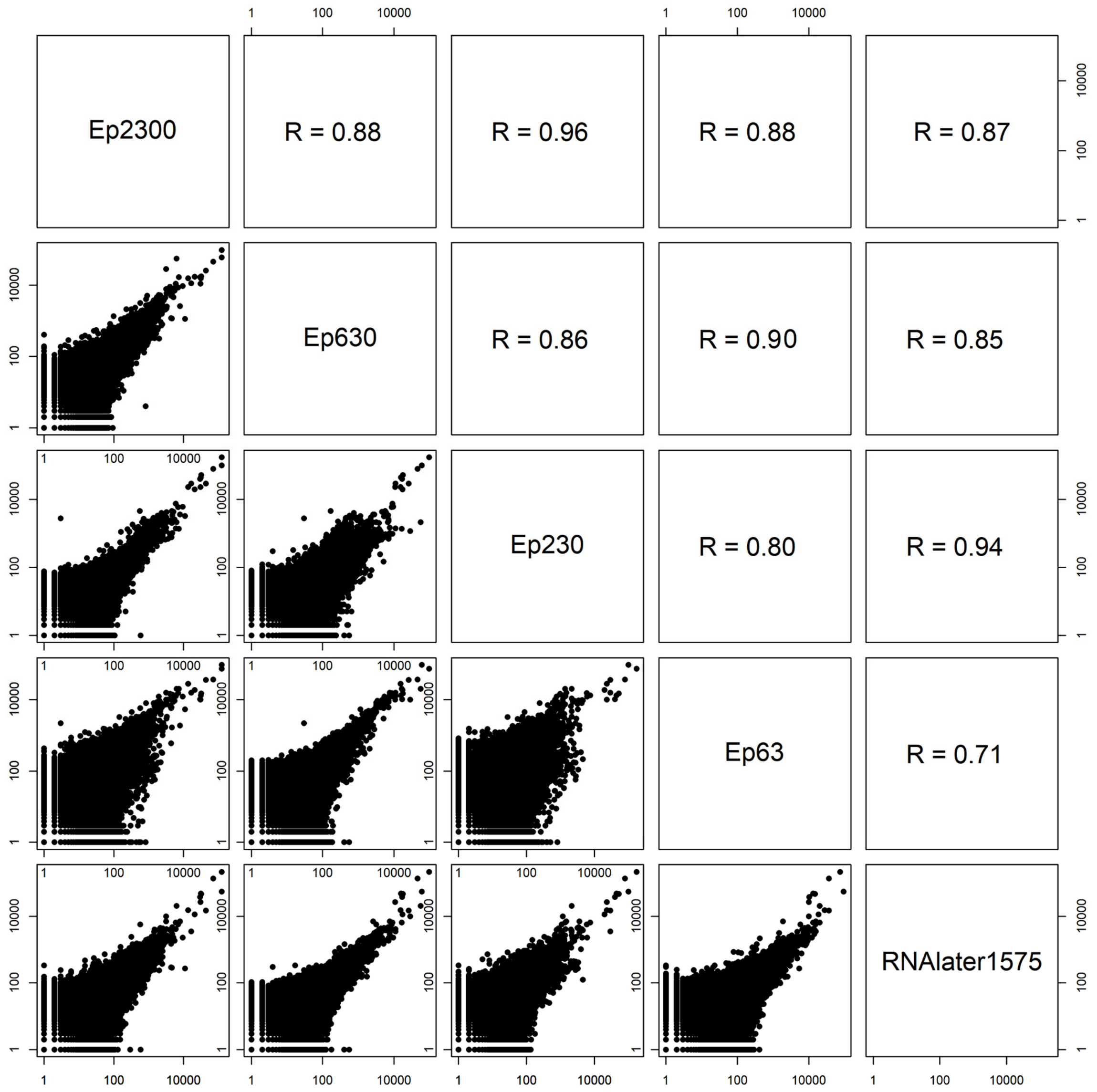


**Figure S12: Reproducibility of the RNAseq data between RNAlater-preserved and snap-frozen preserved mouse intestine sections.** Log-log scatter plots and Pearson correlation coefficients between the RNA data from the ~1575 Ep cells that were collected from the RNAlater-preserved sections and that obtained from the snap-frozen preserved sections. All these data were obtained from the crypt region in the mouse small intestine epithelial cells by immuno-LCM-RNAseq.

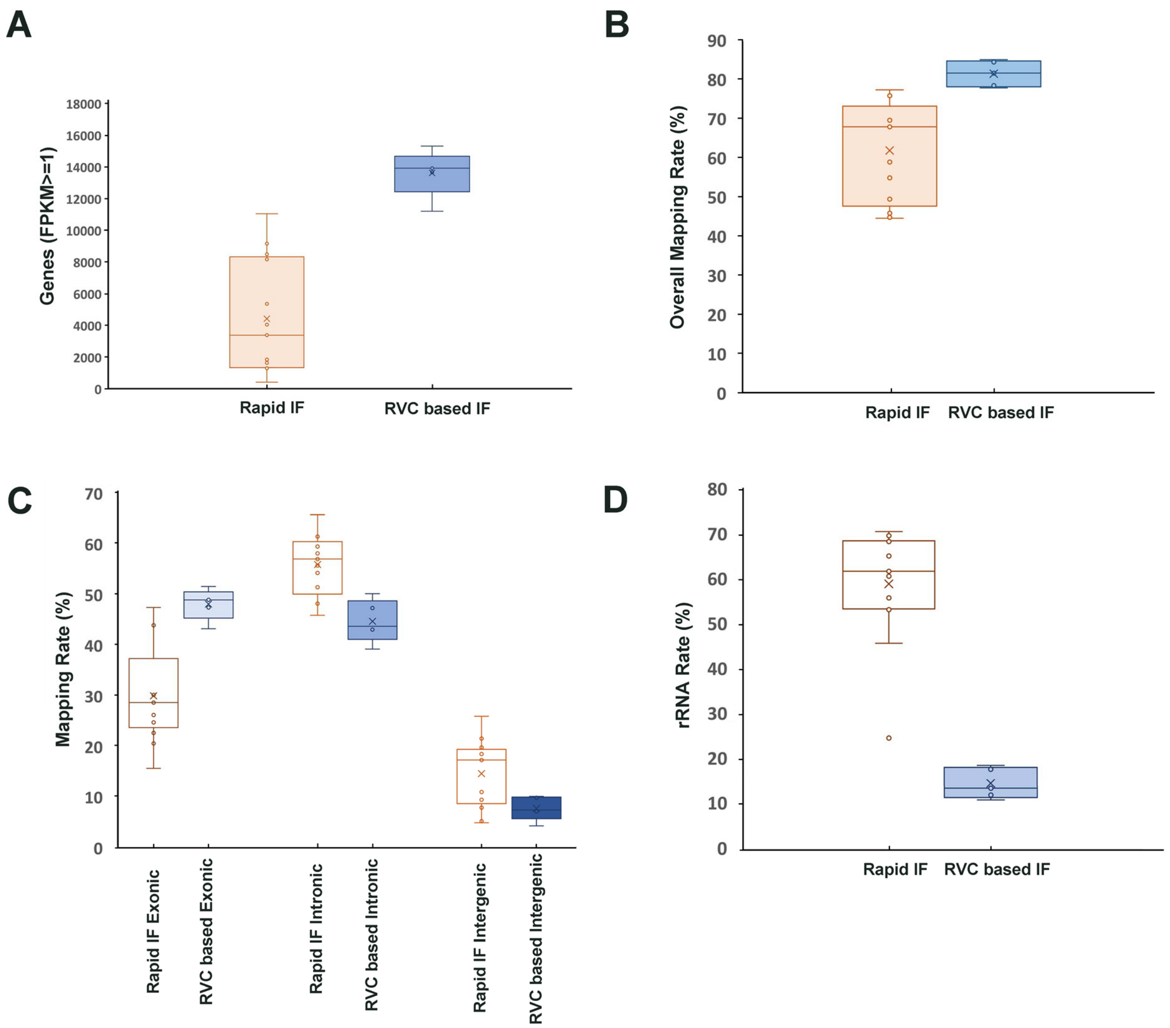


**Figure S13: Comparison of the RNAseq data obtained with our method (small intestine epithelial cells) with that obtained recently using the Rapid protocol** (Baccin et al. 2020)**.** Boxplots show the differences in (A) overall number of detected genes, (B) overall mapping rate, (C) exonic, intronic, and intergenic mapping rate, and (D) rRNA mapping rate between the two approaches. The analysis derived from the data in (Baccin et al. 2020) are from 13 different samples, 9 with a similar read depth as our samples and 4 with 50% greater read depth.

**Table S1**: Number of cells from PanCK-labelled epithelial cells (Ep) in the snap-frozen or RNAlater-preserved mouse intestine samples.

| Samples | Total area ($\mu m^2$) | Effective areal fraction | Section thickness (µm) | Estimated cells number |
|---|---|---|---|---|
| Ep63 | 10800 | 0.68 | 12 | 63 cells |
| Ep230 | 40758 | 0.68 | 12 | 230 cells |
| Ep630 | 108000 | 0.68 | 12 | 630 cells |
| Ep2300 | 407580 | 0.68 | 12 | 2300 cells |
| RNAlater1575 | 270220 | 0.68 | 12 | 1575 cells |

The average volume of intestine epithelial cells was calculated to be 1400 $\mu m^3$.

Cellular areal fraction refers to the fraction of the dissected region that is occupied by cells.

Estimated cells number = Total area * Effective areal fraction * Section thickness /1400 $\mu m^3$

**Table S2**: RNA sequencing information for all of our samples and 13 samples from a recently published Rapid Immunostaining-based LCM RNAseq study (Baccin et al. 2020).

| Samples | Sequencing Clean Reads | rRNA Rate | Overall Mapped Rate | Exonic Rate | Intronic Rate | Intergenic Rate | Genes (FPKM ≥1) |
|---|---|---|---|---|---|---|---|
| Ep63 | 20.9M | 12.00% | 84.90% | 48.77% | 47.14% | 4.08% | 11201 |
| Ep230 | 13.3M | 17.80% | 77.70% | 51.39% | 39.02% | 9.59% | 13657 |
| Ep630 | 19.8M | 10.90% | 84.30% | 43.07% | 50.00% | 6.94% | 15315 |
| Ep2300 | 15.6M | 13.50% | 78.30% | 49.25% | 43.54% | 7.21% | 13913 |
| RNAlater1575 | 15.3M | 18.70% | 81.50% | 47.25% | 42.87% | 9.88% | 14019 |
| H-RNAlater100 | 42.5M | 23.00% | 73.30% | 45.94% | 49.72% | 4.34% | 15819 |
| Lacteal tip and tube samples | | | | | | | |
| Tip_replicate1 | 50.3M | 22.42% | 74.01% | 50.25% | 43.75% | 6.00% | 12389 |
| Tip_replicate2 | 58.3M | 17.77% | 79.22% | 45.77% | 49.88% | 4.35% | 12345 |
| Tip_replicate3 | 55.2M | 16.91% | 80.73% | 47.34% | 48.51% | 4.15% | 12598 |
| Tube_replicate1 | 52.3M | 19.61% | 80.76% | 55.54% | 39.69% | 4.77% | 11771 |
| Tube_replicate2 | 48.7M | 15.43% | 79.10% | 48.18% | 45.32% | 6.49% | 13069 |
| Tube_replicate3 | 62.7M | 15.88% | 77.76% | 42.23% | 51.83% | 5.94% | 14509 |
| Tube_replicate4 | 56.0M | 14.40% | 82.15% | 41.95% | 53.66% | 4.39% | 14223 |
| Bone Marrow – Rapid immunofluorescence staining based LCM RNAseq samples | | | | | | | |
| S1 | 19.5M | 68.82% | 58.83% | 22.99% | 59.27% | 10.73% | 4043 |
| S3 | 22.7M | 55.98% | 76.18% | 43.74% | 48.58% | 7.68% | 9167 |
| S14 | 38.4M | 53.68% | 77.24% | 43.75% | 51.24% | 5.01% | 8487 |
| S15 | 32.2M | 45.90% | 75.74% | 47.26% | 48.04% | 4.70% | 8164 |
| S18 | 18.1M | 68.56% | 44.47% | 24.57% | 54.06% | 21.37% | 1298 |
| S20 | 19.1M | 69.87% | 49.34% | 22.48% | 57.91% | 19.61% | 1348 |
| S22 | 28.2M | 61.91% | 45.78% | 28.49% | 45.74% | 25.77% | 417 |
| S28 | 21.0M | 70.79% | 68.05% | 28.92% | 61.85% | 9.22% | 3379 |
| S29 | 27.5M | 24.79% | 70.41% | 15.52% | 65.56% | 18.92% | 11038 |
| S42 | 21.2M | 68.62% | 67.79% | 30.61% | 58.44% | 10.95% | 1849 |
| S49 | 19.2M | 53.35% | 69.46% | 20.41% | 61.25% | 18.35% | 5362 |
| S75 | 22.7M | 65.34% | 44.70% | 26.04% | 56.81% | 17.15% | 1636 |
| S77 | 20.5M | 60.83% | 54.79% | 26.03% | 55.51% | 18.46% | 1318 |

**Table S3**: Number of cells of Lyve1 labelled lacteal tip and tube samples from snap-frozen- preserved mouse intestine and Podoplanin-labelled lymphatic vessel cells from the RNAlater-preserved clinical human jejunum sample.

| Samples | LCM captured unit | Cells per unit | Section thickness (μm) | Species | Estimated cells number |
|---|---|---|---|---|---|
| **Tip_replicate1** | 101 tips | 1-2 cells | 12 | Mouse | ~150 cells |
| **Tip_replicate2** | 103 tips | 1-2 cells | 12 | Mouse | ~150 cells |
| **Tip_replicate3** | 100 tips | 1-2 cells | 12 | Mouse | ~150 cells |
| **Tube_replicate1** | 28 tubes | 5-7 cells | 12 | Mouse | ~170 cells |
| **Tube_replicate2** | 27 tubes | 5-7 cells | 12 | Mouse | ~170 cells |
| **Tube_replicate3** | 29 tubes | 5-7 cells | 12 | Mouse | ~170 cells |
| **Tube_replicate4** | 28 tubes | 5-7 cells | 12 | Mouse | ~170 cells |
| **H-RNAlater100** | 11 LCM unit | 8-10 cells | 12 | Human | ~ 100 cells |

**Table S4**: Detected isoforms of Arpc1a in the tip and tube samples.

| | **Transcript ID** | **Length (bp)** | **Annotation** | **Expression (FPKM)** | | | | | | |
|---|---|---|---|---|---|---|---|---|---|---|
| | | | | **Tip_ rep1** | **Tip_ rep2** | **Tip_ rep3** | **Tube_ rep1** | **Tube_ rep2** | **Tube_ rep3** | **Tube_ rep4** |
| **Arpc1a-201** | **ENSMUST00000031625** | **1632** | **Protein coding** | **0** | **0** | **0** | **33.43** | **14.63** | **19.08** | **29.10** |
| Arpc1a-202 | ENSMUST00000124379 | 588 | Protein coding | 0 | 0 | 0 | 0 | 0 | 0 | 0 |
| Arpc1a-203 | ENSMUST00000127694 | 1685 | Nonsense mediated decay | 7.48 | 11.28 | 16.62 | 0.38 | 0.73 | 0 | 1.82 |
| Arpc1a-204 | ENSMUST00000134835 | 326 | Processed transcript | 0 | 0 | 0 | 0 | 0 | 0 | 0 |
| Arpc1a-205 | ENSMUST00000142276 | 926 | Retained intron | 15.15 | 3.83 | 10.19 | 0.52 | 6.50 | 6.80 | 2.13 |
| Arpc1a-206 | ENSMUST00000147564 | 425 | Retained intron | 0 | 0 | 1.26 | 0 | 3.27 | 0 | 1.96 |
